# Energy dependence of signalling dynamics and robustness in bacterial two component systems

**DOI:** 10.1101/2023.02.12.528212

**Authors:** Joshua Forrest, Michael Pan, Edmund J. Crampin, Vijay Rajagopal, Michael PH Stumpf

## Abstract

One of the best known ways bacteria cells understand and respond to the environment are through Two-Component Systems (TCS). These signalling systems are highly diverse in function and can detect a range of physical stimuli including molecular concentrations and temperature, with a range of responses including chemotaxis and anaerobic energy production.

TCS exhibit a range of different molecular structures and energy costs, and multiple types co-exist in the same cell. TCSs that incur relatively high energy cost are abundant in biology, despite strong evolutionary pressure to efficiently spend energy.

We are motivated to discern what benefits, if any, the more energetically expensive variants had for a cell.

We seek to answer this question by modelling energy flow through two variants of TCS. This was accomplished using bond graphs, a physics-based modelling framework that accurately models energy transfer through different physical domains. Our analysis demonstrates that energy availability can affect a cell’s signal sensitivity, noise filtering effectiveness, and the stimulus level where cell response is maximal. We also found that these properties are determined not by the molecular parameters themselves, but the reaction rate parameters that govern the reaction systems as a whole.

This suggests possible connections between the molecular structure and evolutionary purpose of any two-component system. This opens the door to new synthetic circuit design in systems biology, and we propose new hypotheses about this link between structure and purpose that could be experimentally verified.

**Author summary:** Two-component systems are the main way many bacteria sense and respond to their environment. They exist in such well-studied bacteria as *E. coli* where they have been shown to detect a range of stimuli including nutrients, temperature, acidity, and pressure.

Two-component systems are ubiquitous in bacteria yet have a deceptively simple structure. Knowing how they operate and the purpose of variations in signalling structure is helpful to our understanding of cellular biology and the design of synthetic biological circuits. Critical unanswered questions remain about the energy usage and functional benefits of these systems.

We sought to improve our understanding of two-component systems by applying a physics-based modelling framework. We found that tracking energy flow through the cell reveals new energy-dependent behaviour in signalling sensitivity, noise filtering, and maximal cell response. We also found that these properties are not strictly dependent on the molecular properties themselves, but from the configuration of the reaction system as a whole.

## Introduction

Two-Component Systems (TCS) are the eyes and ears of bacteria. They are biochemical signalling elements that transmit information from the cellular exterior to the inside of the cell to initiate an appropriate response to environmental cues. TCSs provide the mechanisms by which bacteria sense nutrients, toxins, pH, temperature, and signals from other bacteria [1–11].

The two components of a TCS are a receptor kinase, which typically sits across the cell membrane, and the response regulator, which triggers the appropriate cellular response. TCSs are the primary sensing mechanisms in bacteria but they also exist in other domains of life, especially in plants.

In *Escherichia coli* there are 30 different TCSs that sense and respond to different environmental signals (see Appendix A in S1 Text for a summary table). The TCSs can be divided into two groups characterised by different architectures of the sensor histidine kinase (see Fig. 1). First, the majority of TCSs belong to the orthodox class, where the HK exhibits a single phosphorylation domain; this type accounts for 26 of *E. coli’s* TCSs. Second, the unorthodox class of TCSs has a HK with a more complicated phosphorelay architecture, where a signal, mediated by a phosphate group, travels along three successive HK domains.

**Fig 1.**
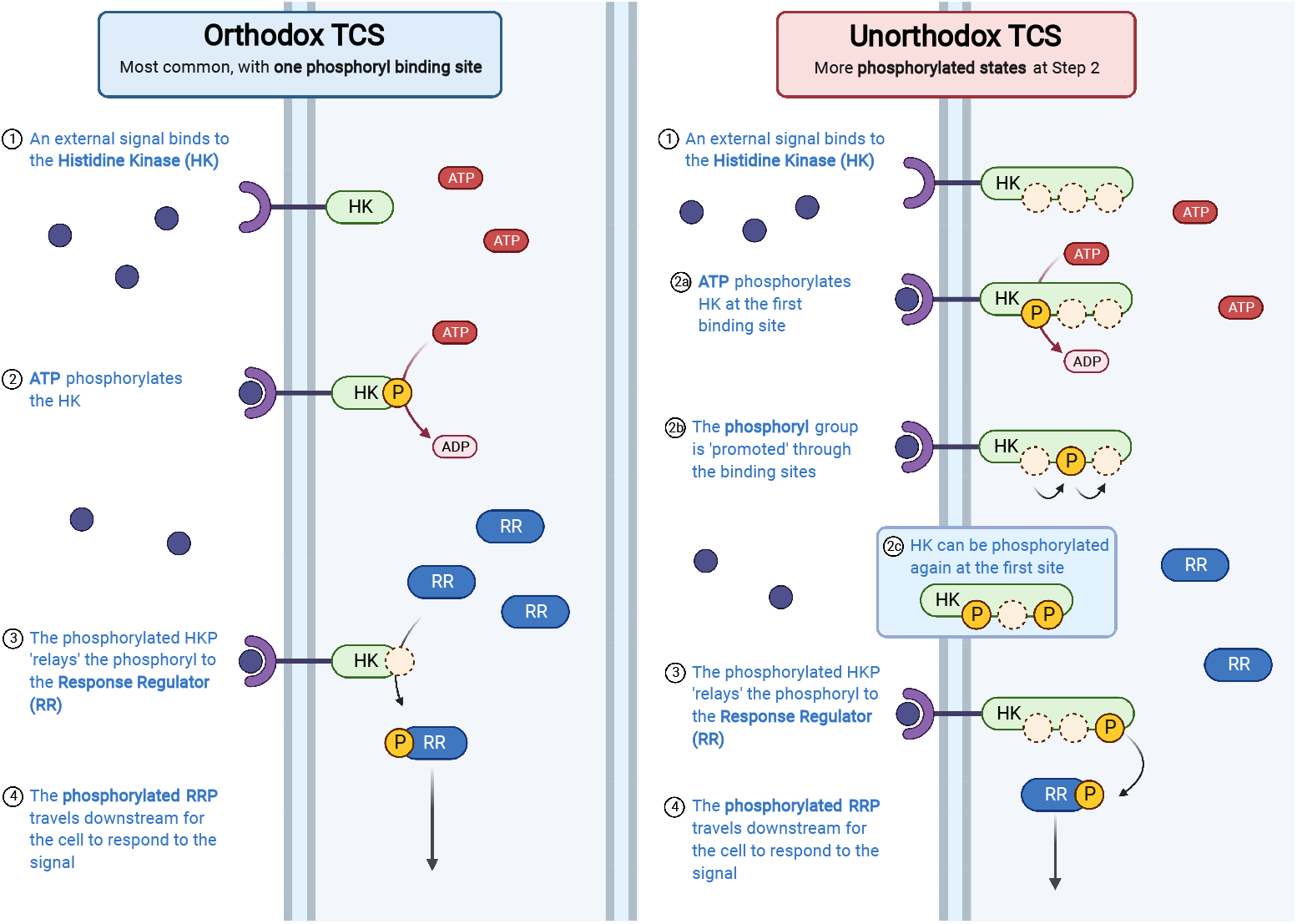
The Two Component Systems (TCS) signalling system found in many bacteria. The two main variants of TCSs shown here are the Orthodox and Unorthodox TCSs. The process in both systems are fundamentally the same: (1) An external stimulus binds or reacts with a Histidine Kinase; (2) ATP phosphorylates the HK; (3) The phosphoryl group is transferred to a Response Regulator; (4) The phosphorylated Response Regulator travels further into the cell to begin a downstream process. Created with BioRender.com.

The reason for the existence of two distinct forms for a sensing mechanism has been a subject of considerable interest and debate. Why do the more complex signalling systems exist alongside the simpler variants? What evolutionary benefits exist for the more elaborate architecture of unorthodox two component systems? Differences in the reliability of information transmission have been put forward. The answers to these questions may help us understand the evolutionary development of bacterial signalling, and how seemingly ‘over-engineered’ reaction systems may actually be crucial for survival.

The amino acid length, and thus energy costs for the different TCSs vary considerably [12]. For example, the EvgSA TCS (involved in drug resistance) requires twice as many amino acids as the QseCB pathway (used for cell motility) and therefore requires more energy to synthesise. All TCS reactions rely on a phosphoryl group supplied by ATP, the primary energy currency of a cell. A TCS with more reactions in its signalling cascade will be energetically more expensive to operate. It is beneficial for a cell to spend ATP efficiently, and yet these energetically costlier TCSs are abundant in biology.

This study aims to address these questions on the qualitative differences between the TCS systems. What different behaviour or functions can TCSs perform? How do TCSs adapt to a changing environment? How well do these systems perform when energy itself is limited - can they function well in low energy (starvation) scenarios?

Mathematical modelling can resolve these issues. A representative model of the dynamics of a class of TCS for arbitrary physical values can tell us more about the structural or variant differences between TCS designs, even synthetic designs [13].

A suite of mathematical modelling studies have already attempted to understand TCS dynamics, their physiological roles, and their evolutionary properties [1,4,7,10,11,14–20]. These models represent the chemical reaction system of TCSs as a set of ordinary differential equations (ODEs) and solve for the response variable numerically, although for some models analytical solutions can be derived [10,11,15,16]. These models have proven useful in investigating the properties of the temporal response to a stimulus and how the response changes as a function of the signal strength [11,14–16,18]. While some modelling studies make use of experimental data [1,4,7], there is often insufficient experimental data to adequately determine the parameter values of these models. Some models address this issue by sampling parameters from a wide range of possible values, allowing one to identify all possible behaviours of a TCS structure [1, 4, 14, 15].

What existing studies lack, however, is an analysis that accounts for the bioenergetics and thermodynamics of signal processing. The dependence on ATP concentration (the energy currency of the cell) is routinely omitted from the models. But TCSs require ATP hydrolysis to drive the signalling cascade. While [14,15] include ATP in their reaction schemes, they assume sufficient ATP for operation and drop it from the model. Accounting for the dependence on ATP concentration is essential for understanding how these signalling systems operate in any environment [21]. We expect that energy abundance in a cell, particularly as expressed in concentration of ATP, will have a measurable impact on the effectiveness and behaviour of cellular subsystems.

TCSs can contain ‘futile cycles’, designs whose function is to dissipate energy. This dissipation can serve important regulatory roles in biochemical pathways, and is important to understand in systems biology models, where ATP (or other energy sources) has to be shared across various cell functions [22, 23].

Here we capture the energetics of molecular signalling in a new mathematical and thermodynamically consistent model of TCS dynamics. Modelling the energetics of biochemical systems has been shown to be useful in predicting realistic behaviour in systems biology [24, 25]. Most biological models enforce some physical laws such as conservation of mass (with e.g. mass action kinetics), but few go as far as to enforce conservation (or realistic dissipation) of energy. Bond graphs are a generalised energy-based modelling framework that provide a modular structure to our system. Bond graphs are a generalised energy-based modelling framework that (i) capture cellular biophysics and thermodynamics; and (ii) provide a modular model structure that is the hallmark of many biological systems. See Methods for more details on bond graphs.

From this energy-based perspective, we create new insights about the purpose and structure of TCSs in different energy environments. In particular, we find that (1) Energy availability affects the performance and sensitivity of signalling systems. (2) Energy availability dictates the signal to noise ratio of TCS signalling. (3) Some systems can filter out noise and eliminate noise propagation in high energy systems. (4) TCS systems can dynamically select particular stimulus levels, in an energy dependent manner..

### Orthodox and Unorthodox Two Component Systems

The most common TCS structures are the orthodox and unorthodox variants [2,4,5,9,26,27]. All variants share the same core signalling structure: a membrane bound histidine kinase (HK) and a response regulator (RR). In the presence of an external stimulus, the HK is phosphorylated on its binding domain. In the orthodox TCS, the HK consists of a single histidine phosphotransfer domain and an ATP binding domain; the unorthodox HK has three binding domains. This phosphoryl group is passed on to the RR either immediately (orthodox) or after a series of phosphorelay reactions (unorthodox) to produce the response species RRP. This RRP is the signalling output which travels into the cell interior to initiate a response to the signal (see Figure 1).

Our study considers how energetics affect TCS signalling and pay particular attention to differences between the two predominant TCS architectures, the orthodox and unorthodox systems. These differ structurally, and the structural differences affect their dynamical characteristics. There has been considerable interest in why these different architectures exist alongside each other. For example, the unorthodox variant has a more complicated reaction system, and the proteins involved are on average longer than their orthodox counterparts (Table 3). In *E. coli*, 4 of 30 TCSs are of the unorthodox type. It has been suggested that evolution may favour different architectures for different signalling scenarios. Modelling studies have shed light on differences between orthodox and unorthodox TCSs: for example, the unorthodox systems appear to be more robust to noise [15]. However, from Table 3, unorthodox TCSs are also more costly to synthesise, which may point towards trade-offs between cost of production and performance.

### Orthodox TCS

The orthodox TCS model is the simplest and most common form of a TCS. The histidine kinase can be in one of two states: unphosphorylated (HK) or phosphorylated (HKP). We model this as a two-step process; HK binds or is otherwise activated by the stimulant (S) to produce HKS. This stimulated species is then autophosphorylated by ATP to produce HKP. HKP then passes on the phosphoryl group to the response regulator (RR) molecule, in a process called phosphorelay to produce a phosphorylated response regulator (RRP). RRP then travels into the cell interior to trigger a downstream process. Eventually, RRP decays back into RR. The chemical reactions for this process are summarised in Table 4.

### Unorthodox TCS

The unorthodox TCS is a more complex variant of the TCS in which the HK goes through a three-step process to phosphorylate the RR. In this variant, there are three distinct binding sites on an HK available for a phosphoryl group. This means that an HK can be in one of 2^3^ = 8 phosphorylated states. We have labelled these HK1 through HK8 - see Fig 2 for the names of all HK states.

**Fig 2.**
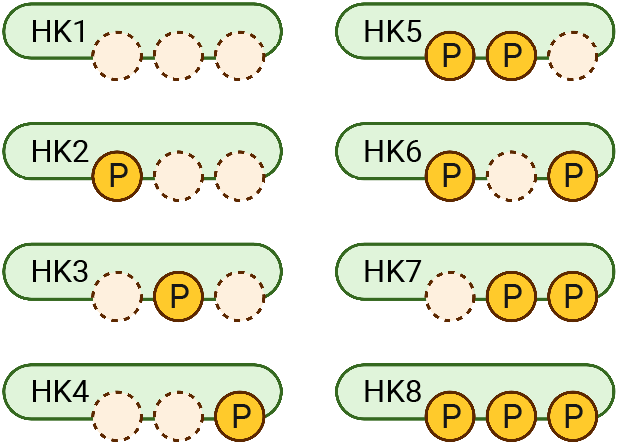
All HK phosphorylation states in our model of the unorthodox TCS. The first binding site is the start of the signalling cascade, and the last binding site is received by the response regulator. HK can move between different state non-sequentially (see Table 2). Naming convention is the same as in [15]. Created with BioRender.com.

There are now multiple parallel autophosphorylation reactions for the unorthodox system. Four of these states (HK1, 3, 4 and 7), which have the first binding site empty, respond to the stimulus and act as starting points for the signalling process. As these phosphorylated HK collide with each other, they promote their phosphoryl group to a higher binding site through a forward phosphorelay. The four states with the final site bound (HK4, 6, 7 and 8) can donate their phosphoryl group to the RR molecule. Each of the above reactions can also spontaneously unbind (i.e. reverse). Eventually, RRP decays back into its original state. The chemical reaction network for this process is summarised in Table 6. As we will discuss later, the multiple pathways that lead to the phospohrylation of RR allow for extra complexity and adaptive behaviour by the TCS.

## Results

We investigate the role of energetics on TCS signalling through mathematical modelling with bond graphs. As discussed earlier there are limited kinetic data available for TCSs. Accordingly, we have simulated many instances of this model with different parameters to cover the broad range of physiological possibilities and explore all the dynamics possible in TCSs. Unlike previous studies, we treat ATP concentration as a tunable parameter. Details on model construction, parameter sampling and simulation protocols are given in Methods. See Appendix B in S1 Text for the full mathematical description.

### ATP increases maximum signal response

To investigate how ATP affects signalling strength, we ran the orthodox and unorthodox models over 10,000 simulations and plotted the response against time (Fig 3). The RRP Fraction is defined as the amount of phosphorylated RR (RRP) normalised by the theoretical maximum RRP expression attained with unlimited ATP. We represent the RRP Fraction as a temporal response curve, which is also called the RRP curve. The response curves for the orthodox and unorthodox TCS models are shown in Fig 3.

**Fig 3.**
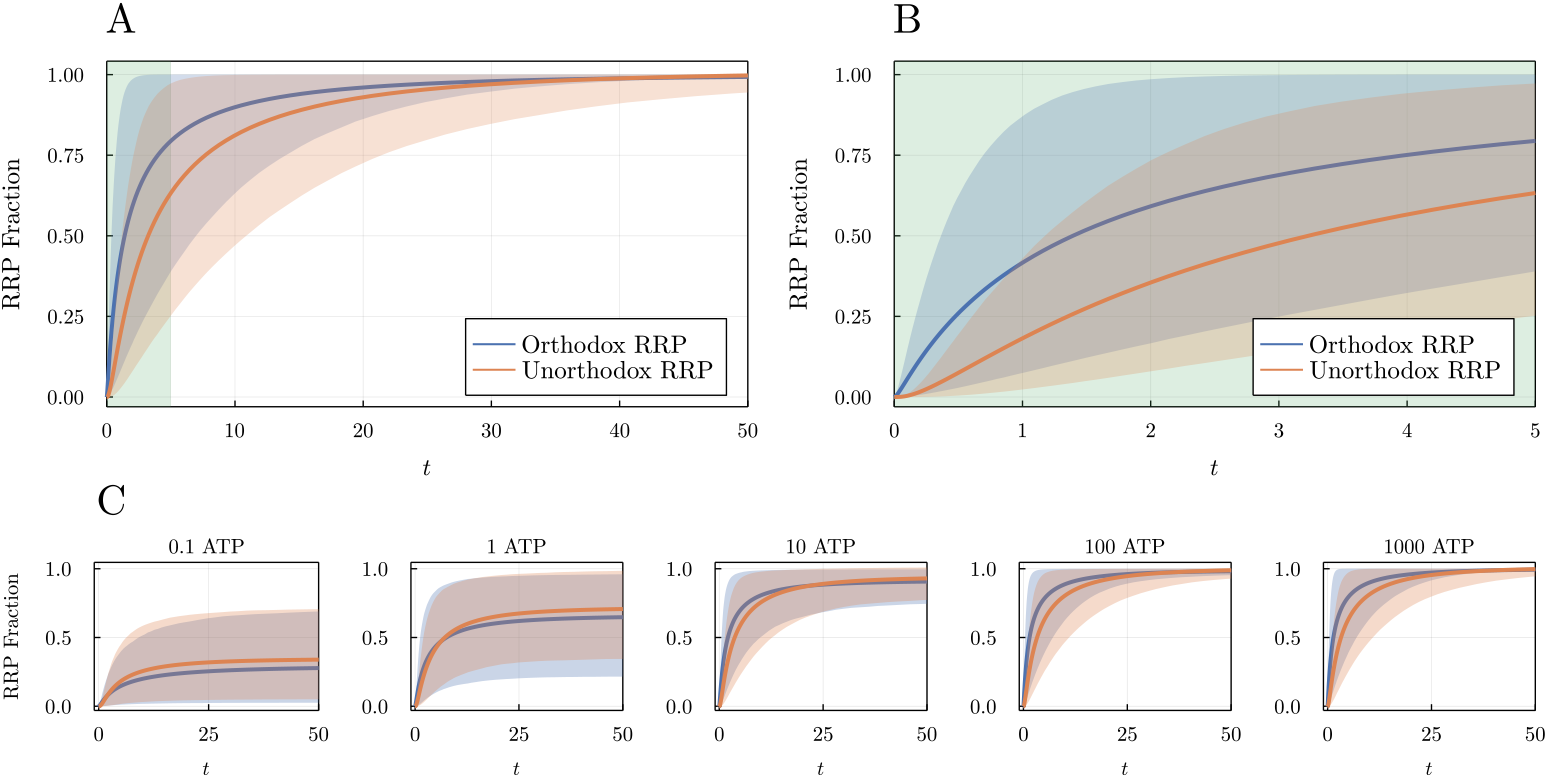
RRP fraction over time after the presence of a signal at *t* =0 for the orthodox (blue) and unorthodox (orange) TCS models. **(A)** The RRP curves are each normalised by their maximum expression value and then averaged over 10,000 simulations with parameters sampled from a log-uniform distribution from 0.1 to 10 (9 parameters for orthodox, 35 for unorthodox). The shaded regions capture the RRP curves between the 10th and 90th percentile (ranked by expression level). **(B)** Zoomed in plot of figure (A) for t **≤** 5, as indicated by the green band. (B) highlights that the orthodox curve is more hyperbolic, and the unorthodox curve is more sigmoidal. **(C)** RRP curves for each TCS model at increasing available ATP levels. Less ATP availability results in a lower signal strength and a slower response, but the maximum signal strength and speed saturates after sufficient energy level. ATP levels are unitless.

On average, the temporal response curves are consistent with earlier studies [15]; the averaged orthodox response curve appears hyperbolic, while the averaged unorthodox response curve appears more sigmoidal (Fig 3A, solid lines). In Fig 3B, we focus on the first 5 seconds of Fig 3A where this difference is more distinct; the unorthodox response curve takes slightly longer to reach maximum signal and resembles a sigmoidal Hill function.

We found that the maximum expression level of this temporal response curve depends on the amount of ATP available. With higher ATP, a higher maximum response signal is possible (Fig 3C). When ATP concentration is low (leftmost subplot), the maximum response is (on average) less than 50% of value at the highest energy level. The highest response is reached at about 100 ATP before the effect saturates. This is true for both the orthodox and unorthodox models. A TCS in a low energy environment may not be able to respond to external stimuli as effectively as in an energy rich environment.

We find here that the average behaviour of the TCS variants aligns with what may expect from previous studies. What is apparent from Fig 3 however is that there is a wide band of possible response curve behaviours for a given instance of a TCS. As we will next see, the distinction between an orthodox and unorthodox TCSs response curve may be hard to identify.

### ATP affects sensitivity

We next investigated how ATP affects the signalling sensitivity of a TCS. We can quantify the sensitivity of a response by calculating the Hill coefficient, which can be estimated using

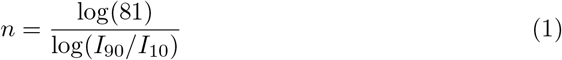

where *I*_90_, *I*_10_ are the concentrations of the stimulus when the response is at 90% and 10% of the maximum response steady state value respectively [15,28]. A coefficient of *n* ≈ 1 corresponds to a subsensitive signal, whereas a coefficient of *n* > 1 corresponds to an supersensitive signal.

The temporal sensitivity of a signal determines how quickly or slowly the signal responds to a stimulus: a subsensitive curve (Fig 4C, top) resembles a hyperbolic curve with a rapid initial response expression; an supersensitive curve (Fig 4C, bottom) resembles the characteristic sigmoidal Hill function, with a slower uptake followed by a rapid increase once the stimulus duration has crossed a threshold.

**Fig 4.**
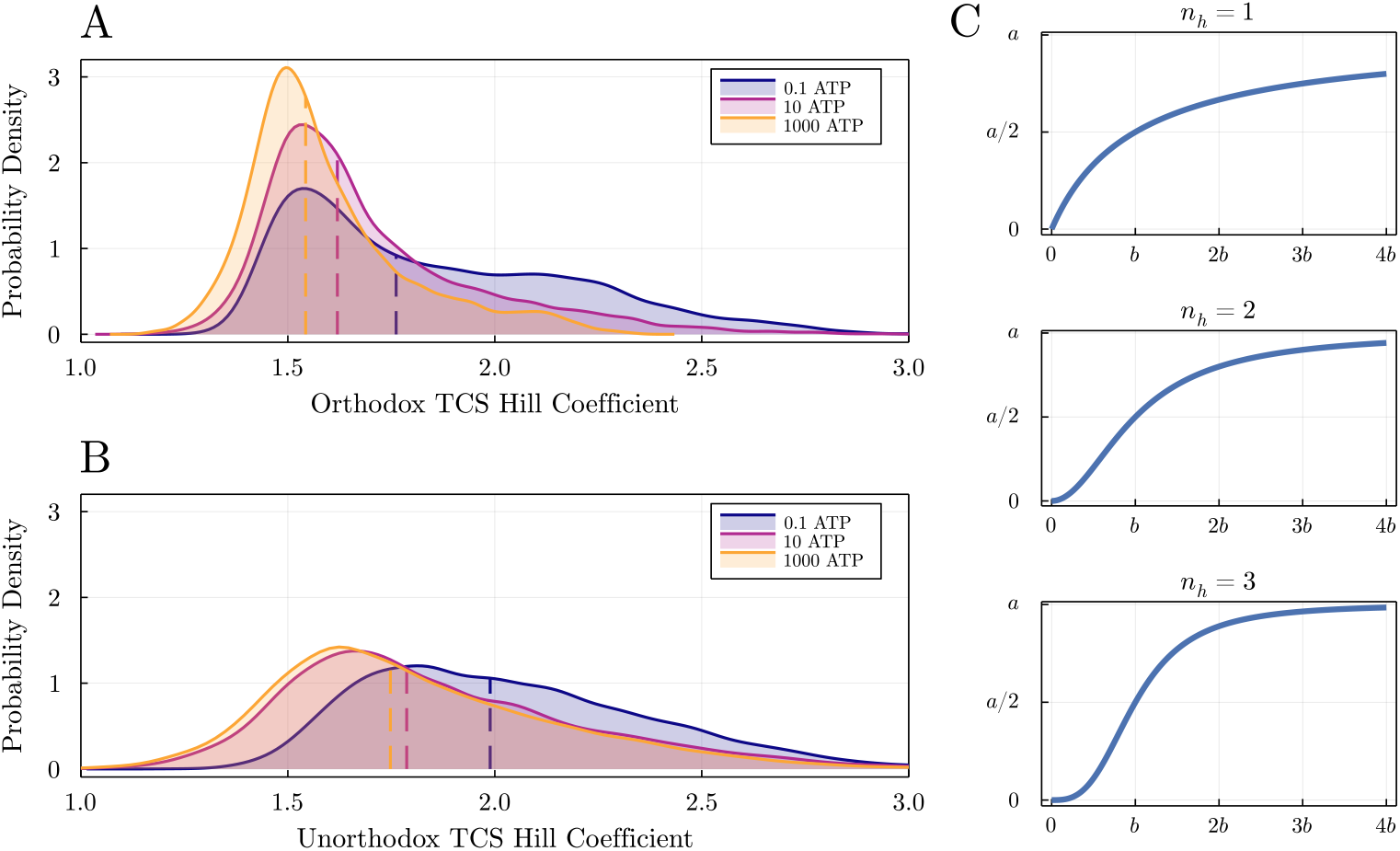
Hill coefficient distributions for **(A)** Orthodox TCS and **(B)** Unorthodox TCS. Dotted lines indicate median values for each distribution. The Hill coefficient is a measure of sensitivity and is defined by the Hill function 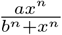, where *n* is the Hill coefficient. It can be estimated using 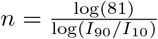, where *I*_90_, *I*_10_ are the concentrations of the stimulus when the response is at 90% and 10% of the maximum RRP steady state value respectively [15,28]. **(C)** Hill functions for *n_h_* = 1, 2, 3. As the Hill coefficient increases, the response curves becomes increasingly sigmoidal.

In Fig 3B, we see that the unorthodox TCS is more sensitive than the orthodox TCS, which aligns with previous modelling studies [4, 15]. However, this simple dichotomy does not capture the true range of dynamics displayed by the systems. Fig 4A and 4B plot the distribution of estimated Hill coefficients for the 10,000 simulations of both TCS models, at different levels of ATP concentration. There are orthodox systems with *n* > 2 and unorthodox systems with *n* < 1.5.

We also see a shift in the distributions to the right as ATP levels decrease (Fig 3A,B). This implies that when ATP availability is reduced, a signalling system will have a higher Hill coefficient and becomes more sensitive to stimuli. This is most striking for the orthodox TCS model (Fig 4A). For higher ATP levels in the orthodox model, the median Hill coefficient is around 1.54. As ATP decreases, the distribution flattens out towards the right, where we see a higher proportion of Hill coefficients above 2 and even higher than 2.5. The system moves from behaving as semi-hyperbolic to behaving like a sigmoidal switch. Under these low energy conditions, an orthodox TCS will appear to have the same sensitive behaviour as an unorthodox TCS.

This shift towards sigmoidal behaviour is less pronounced in the unorthodox TCS model, as this model tends to already exhibit supersensitive behaviour. This distribution remains stable even for low energy levels (10 ATP), but does still shift for extremely low ATP concentrations (0.1 ATP).

### ATP determines noise filtering effectiveness

So far we have only considered signalling behaviour for an ideal stimulus: constant, stable, and unambiguous. We next look at signalling performance in the presence of a noisy stimulus; that is, a stimulus detected with an unbiased error (see Methods for mathematical details). A noisy stimulus may represent uncertainty in detecting the stimulus (especially at low concentrations), but it may also be an erroneous signal that the cell should not respond to. It is important therefore that critical signalling systems are able to filter out noise, or at least reduce its impact.

Orthodox and unorthodox variants respond to noise differently. Specifically, the unorthodox TCS structure is able to filter erroneous noise more effectively than the orthodox TCS [4,14,15]. We investigate this phenomenon using our bond graph model and suggest why this occurs. We repeat earlier simulations using the same generated parameter values but with a noisy stimulus input. We then compare the same TCS instance response with both a constant and a noisy signal.

We find that unorthodox TCS instances filter noise more effectively than orthodox TCS instances (Figs 5A, B). For the same noisy stimulus input, the averaged response curves for the orthodox models (5A) exhibit more frequent spikes in the signal, when compared to the equivalent response for the unorthodox models (5B). The unorthodox TCS model appears to be better at handling noisy inputs on average, in line with previous studies. Moreover, the variability in the output is less prominent at high energy levels, compared to the orthodox system.

**Fig 5.**
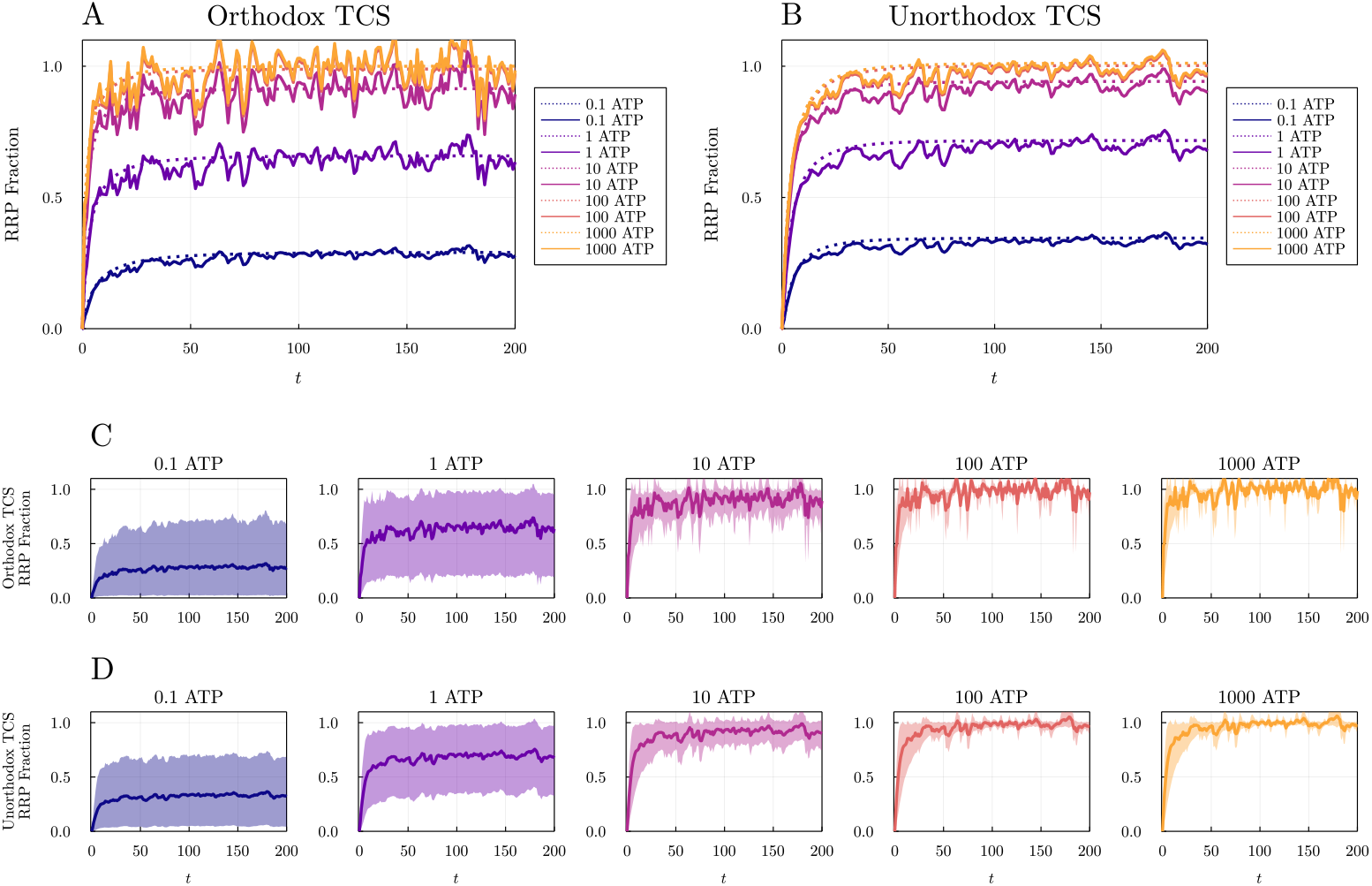
Temporal response curves (RRP Fraction) for TCS models when exposed to a noisy stimulus. The response curves for each ATP level averaged over all parameter set simulations for the **(A)** orthodox and **(B)** unorthodox TCS models. The response curves for a noisy stimulus (solid line) are compared to the response from a constant stimulus (dotted line). The averaged noisy response curves for **(C)** orthodox and **(D)** unorthodox TCSs over the same simulations are shown with 10-90th percentile ribbons.

When the aggregated response curves for a noisy signal (5A, B, solid lines) are plotted against the aggregated response curves for a constant signal (5A, B, dashed lines), we see that the noisy signals tend to give a slightly lower response level compared to their constant counterparts. This means that the mean signal magnitude is reduced by the extra noise; a noisy stimulus produces a slightly weaker signal strength. This appears to be the case on average across all models and energy levels.

Additionally, we find an interesting effect of energy availability on noise filtering. As ATP concentration is increased, the orthodox TCS amplifies the magnitude of the noise. With higher energy availability, the magnitude and variance of the noisy response orthodox curve increases (5C). On the other hand, the unorthodox TCS models are much better at noise filtering, even at the highest energy levels (5D).

We compared noise filtering performance with three simple measures: the signal mean *μ*, standard deviation *σ*, and coefficient of variation *σ/μ* (Fig 6). The coefficient of variation was used to compare the variability (standard deviation) of different distributions. These measures were calculated for the RRP concentration for each simulation run, each model structure, and each level of ATP.

**Fig 6.**
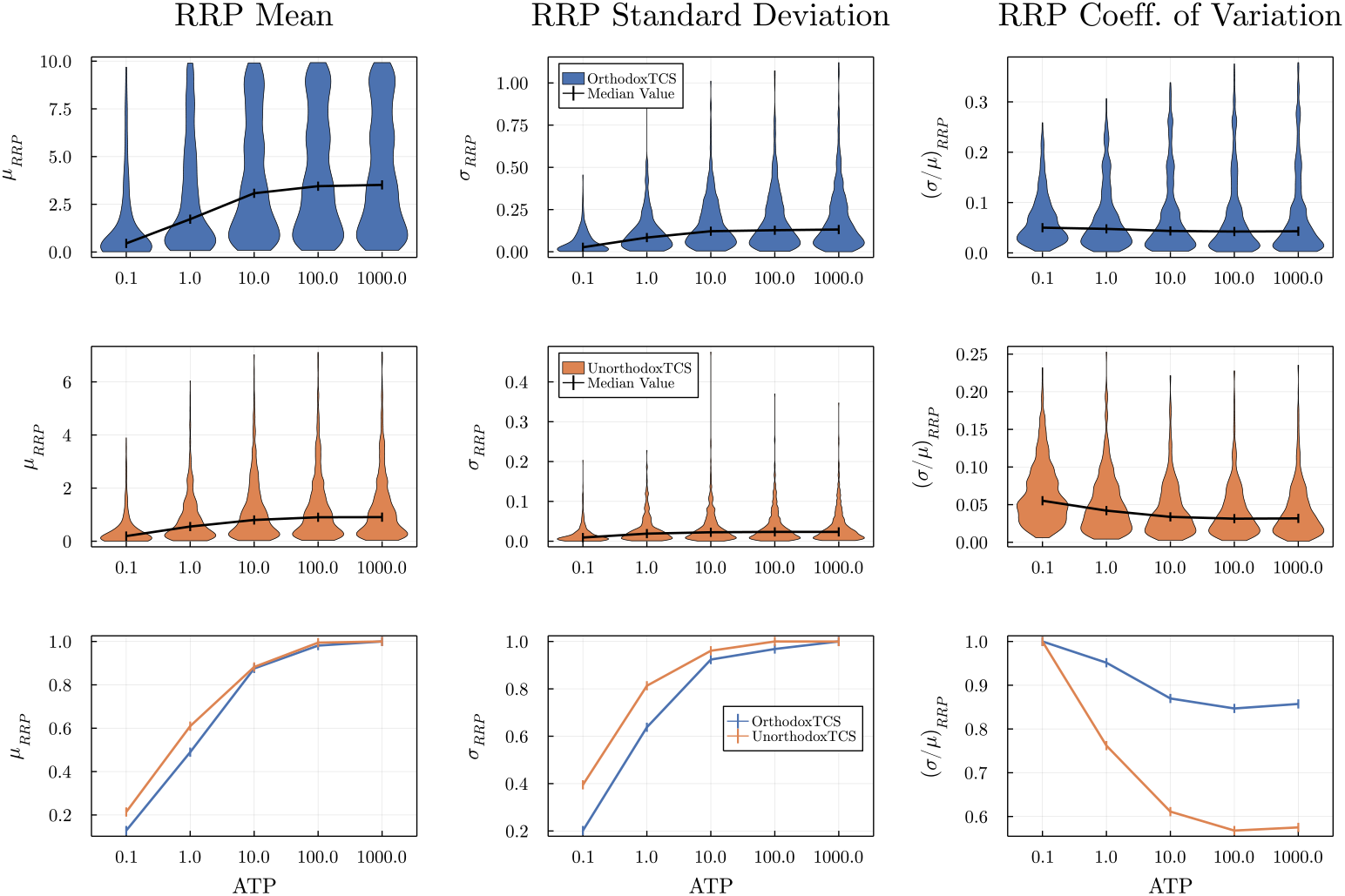
Performance metrics for the noisy response curves as ATP concentration increases for orthodox (blue) and unorthodox (orange) TCSs. The mean *μ*, standard deviation *σ*, and coefficient of variation *σ/μ* of the RRP concentration were calculated after a warm-up period of 100s. The last row of plots compares the normalised median values of both TCS models for each metric.

For both TCS models, the median signal response drops as ATP levels decrease (Fig 6, left column). This is in agreement with what we see in Figs 3C and 3D. The standard deviation of the signal for orthodox TCSs increases with more energy, while it remains stable for the unorthodox system (Fig 6, middle column). Again, this follows from what we have seen in Fig 5. However, the median coefficient of variation for the orthodox systems does seem to decrease with more ATP, although this decrease in variation is less than for the unorthodox systems (Fig 6, right column). The unorthodox TCSs are therefore on average more effective at reducing noise in the signal response.

Why does the unorthodox TCS filter noise better than an orthodox TCS? We begin to see why this may be by examining the reaction fluxes for each model. The bond graph framework allows us to extract the fluxes over time for each reaction in our model. Fig 7 plots the reaction fluxes for two exemplar instances of an orthodox TCS (top) and unorthodox TCS (bottom), along with their response curves. The reaction fluxes (black) are overlaid on top of each other to highlight the cumulative effect of the whole system.

**Fig 7.**
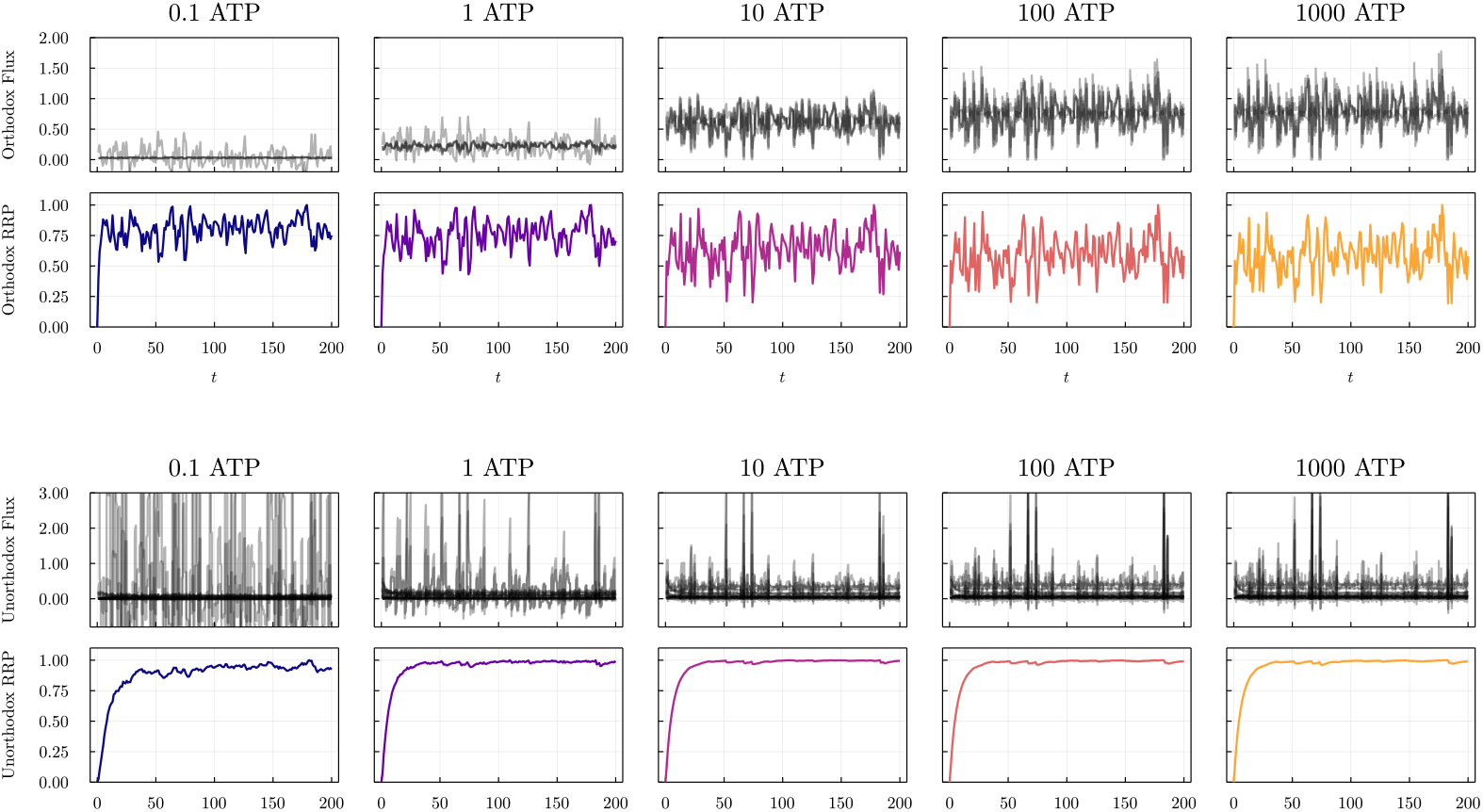
Comparing reaction fluxes of all reactions in the TCSs for example orthodox TCSs (top two rows) and unorthodox TCSs (bottom two rows) simulations. ATP levels increase across the figures from left to right. The upper plots are the flux for each reaction in the model as outlined in Table 2, and are overlaid over each other in black. The lower plots are the corresponding response curve outputs. For the orthodox model, increasing ATP not only increases baseline flux and flux variability, the apparent baseline signal of the response curve also decreases; more ATP leads to a noisier, less reliable signal despite the higher reaction throughput. The unorthodox model however is robust to additional ATP; the noise is dampened, the signal is maintained, and the reaction flux does not vary considerably.

Fig 7 gives us three interesting observations. For the orthodox TCS, increasing ATP has three effects: (1) increases the baseline level of reaction flux, meaning each reaction is happening more often; (2) it increases the variability (noise) of the flux; and (3) it the baseline level for the response curve (second row) decreases as the signal becomes noisier. More ATP leads to a noisier, less reliable signal despite the higher reaction throughput. By contrast, the unorthodox TCS here does not have the same trends. In fact, the flux variability decreases with additional ATP (third row). The response curve (fourth row) is robust to additional ATP; the noise is dampened, the signal is maintained, and the reaction flux does not vary considerably.

### Unorthodox TCS selects for specific stimulus ranges

We finally examine the Signal-Response (SR) curve for the TCS variants. SR curves show the relationship between the stimulus and the response at steady state. We repeat simulations for a wide range of stimulus levels and determined the final steady state response expression for each run.

The SR curve for all the orthodox TCS we investigate resemble monotonic logistic curves (Fig 8, top row). This aligns with what we expect from existing models of TCSs [14–16]. However, for the unorthodox TCSs, we find that only 22% of the SR curves are monotonic at every ATP concentration (Table 1a). In the non-monotonic cases, the maximum response expression level is not found at maximum stimulus expression (Fig 8, bottom row). This means that a system which optimises for a maximum signal response expression will select for an intermediate stimulus value, not the maximum possible value. Note that only unorthodox systems have this property; no orthodox system studied had a non-monotonic SR curve.

**Fig 8.**
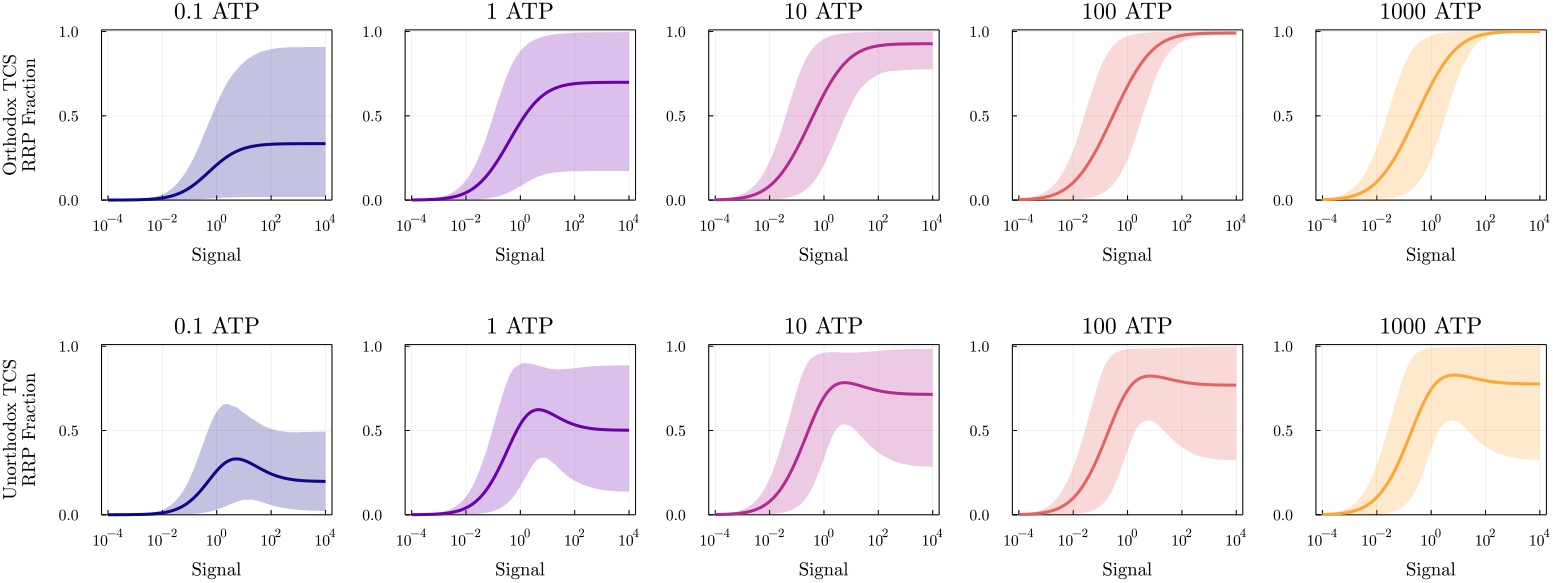
Signal Response (SR) curve plots for the orthodox and unorthodox TCS models for increasing ATP concentrations. SR plots show the final RRP steady state value (as a fraction of the maximum RRP value) for a range of stimulus sizes (plotted on a log scale). The solid line is the mean SR curve over 10,000 parameter sets and the ribbon is the 10-90th percentile range. The orthodox SR curves are all sigmoidal, whereas for the unorthodox SR curves can exhibit non-monotonic behaviour.

**Table 1.** Proportions of SR Curve Classifications for Unorthodox TCS. Proportions of SR curve classes by shape and peak shift for the unorthodox TCS simulations. Examples from each class are shown in Fig. 10 and Fig. 11. The Peak Shift classes are only defined for the SR curves which are All Non Monotonic.

| Class | Count |
| --- | --- |
| All Monotonic | 2206 |
| Peak Lost | 2409 |
| Peak Gained | 669 |
| All Non Monotonic | 4716 |
| Total | 10000 |

(a) SR Curve Shape
| SR Curve Peak Shift | Count |
| --- | --- |
| Right Shift | 381 |
| No Shift | 1414 |
| Left Shift | 2921 |
| Total | 4716 |

(b) SR Curve Peak Shift

Remarkably, the existence of a non-monotonic peak in an SR curve is itself energy dependent. For some unorthodox TCSs, the SR curve is only monotonic at low energy levels, and non-monotonic when energy levels are increased. For other unorthodox TCSs, the opposite is true: the curve is non-monotonic at low energy, and monotonic at high energy. We have categorised the SR curve shapes into four classes (Fig. 10):

1. **All Monotonic** Monotonic SR curves at all energy levels
2. **Peak Gained** Non-monotonic at higher energy levels only
3. **Peak Lost** Non-monotonic at low energy levels only
4. **All Non Monotonic** Non-monotonic SR curves at all energy levels.

These classes can be grouped together in several ways. In a low-energy environment (<0.1 ATP), the All Monotonic and Peak Gained classes are similar, and the All Non Monotonic and Peak Lost are similar. In a high energy environment (>100 ATP), the All Monotonic and Peak Lost classes are similar, and the All Non Monotonic and Peak Gained are similar. Roughly 30% of the unorthodox TCSs instances switched their behaviour across the ATP gradient. Surprisingly, of the 10,000 simulations we ran, we found that only 22% of unorthodox TCSs had monotonic SR curves at all energy levels.

Similarly, we also find that the position of the SR curve peak can also move across the ATP gradient. The stimulus required for the optimal signal response may change for different energy levels. The peak of the signalling curve can shift to the left or right as energy levels increase (e.g. the All Non Monotonic example in Fig 10). We have likewise divided the non monotonic unorthodox TCSs into peak-shift classes (Fig 11): A **Right Shift** means that as ATP increases, the signal peak moves to the right, thereby maximising the response for a higher signal strength (about 8% of cases). A **Left Shift** means that as energy increases, the response is maximised at a lower signal level (about 62% of cases). In 30% of simulations, there is no significant movement in one direction across energy levels (**No Shift**), though the peak may move slightly between levels. Note that these classes are only defined where a peak exists at all energy levels (i.e. only All Non Monotonic SR curves).

In some rare cases (<0.5% of simulations) the solution exhibited ‘multi-peak’ behaviour, where there are two or more local maxima in the SR curve. It is currently unclear what leads to this phenomenon.

At this stage it is difficult to predict which class a solution will fit in from the parameter values alone. To draw some insight we began exploratory analysis of the distribution of parameter values and which SR curves they tended to create.

We first looked at how the parameter values were distributed for the monotonic behaviour of the SR curves. We plot the distribution of the 21 unorthodox reaction rate parameters *κ* in Fig. 9, split by our SR classes (see S1 Text for a similar plot of the 14 thermodynamic constants *K*). From these plots we can already make several observations.

**Fig 9.**
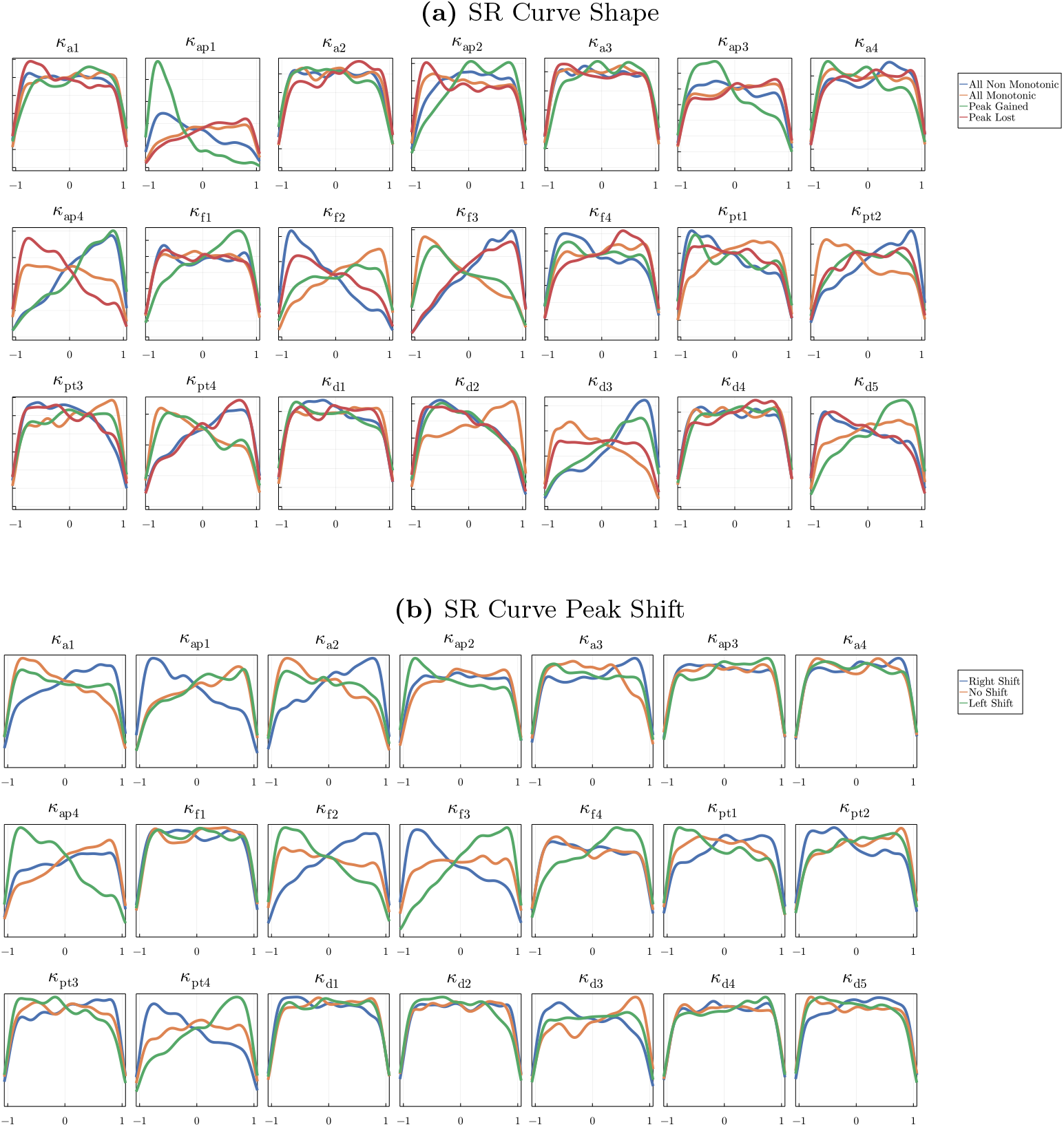
Log distribution of the sampled reaction rates *κ* used for the unorthodox TCSs numerical simulations, split by **(a)** SR curve shape and **(b)** SR peak shift (see Figs. 10 and 11). The reaction rate variables are the same as in Table 2 and defined in Appendix B in S1 Text. Parameters are sampled from a log-uniform distribution (Eq. 8). Without any group effects we would expect all classes to follow a uniform distribution on these log plots; however particular reaction rate values bias towards particular SR curve shapes. For example, a low value for *κ*_ap1_ (row 1, column 2) biases heavily towards the Peak Gained and All Non Monotonic shape classes, and the Right Shift peak-shift class. By contrast, the distributions for the thermodynamic constants *K* (Appendix C in S1 Text) all form the same distribution, and so do not determine SR curve (energy dependent) behaviour.

First, the difference in behaviour is determined entirely by the reaction rates *κ*. The distributions for the reaction rates all take different shapes for each class; in contrast, the thermodynamic constants K follow the same distribution shape regardless of class (Appendix C in S1 Text). The reaction rates are what determine these SR curve behaviours. Since the thermodynamic constants are inherent properties of the species, we see that this dynamic behaviour is not a result of species but of the reaction network itself. Exotic behaviour cannot be identified by analysing the chemical species alone; it is necessary to consider the network structure in its entirety.

Second, certain reaction rates stand out as having greater explanatory power. For example, when *κ*_ap1_ (row 1, column 2) is low, we are more likely to get a Non Monotonic (blue) or Peak Gained (green) solution. This parameter is responsible for the initial rate of phosphorylation of the HK, so we can infer that slow binding of ATP to the HK domain can cause peaks in the SR curve. A faster binding rate for ATP results in a monotonic switch for the SR. To get the signal selecting behaviour, this early stage reaction should be slow relative to the other TCS reactions. Incidentally, these classes resemble each other at high energy levels (bottom row of Fig 10).

**Fig 10.**
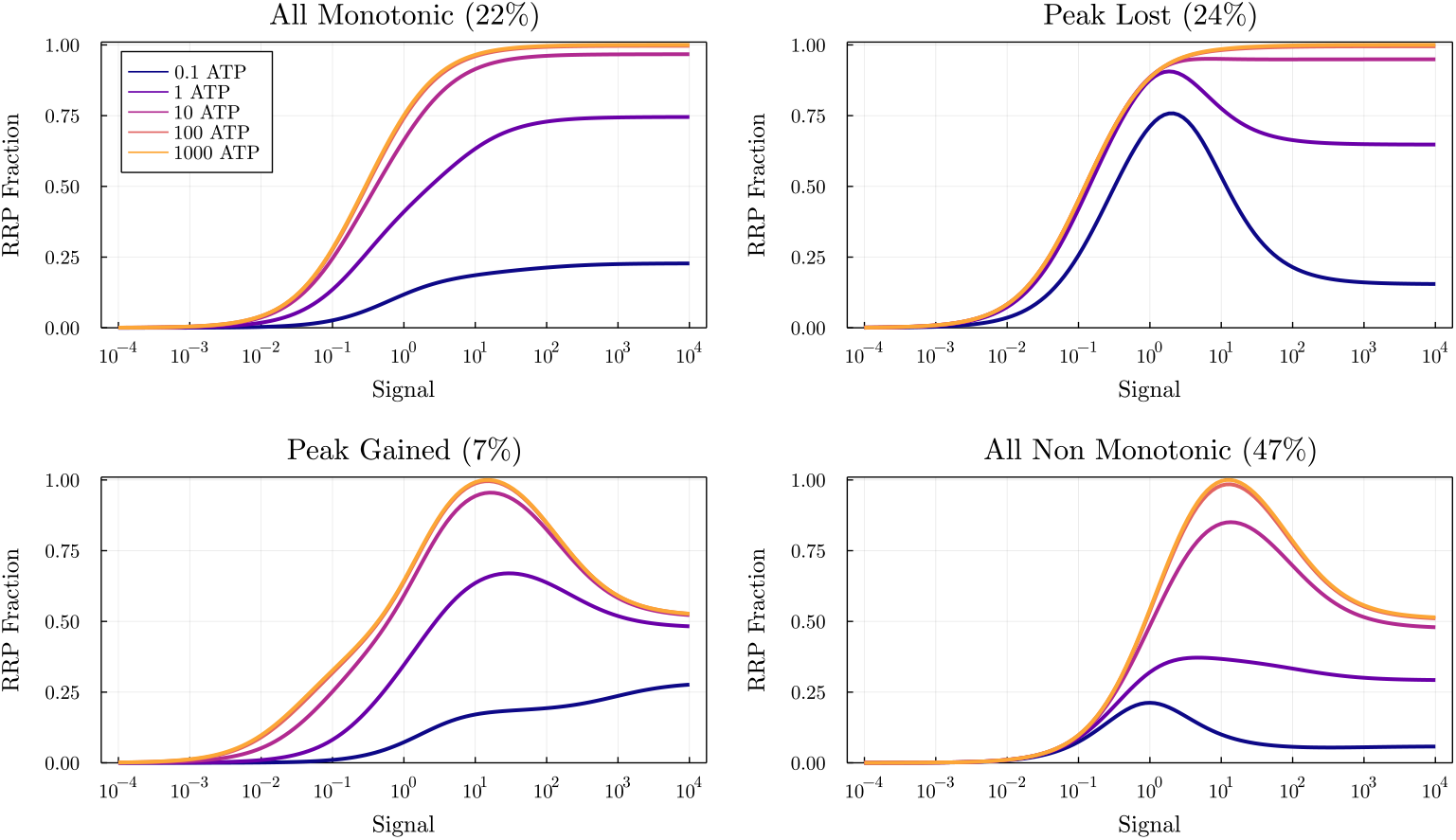
Representative examples of the SR curve shape classes for the unorthodox TCS simulations. The proportions of each class are included in parentheses. These classes partition each simulation instance based on how the SR curve shape changes as ATP is increased or decreased, all else being equal. **All Monotonic** Monotonic SR curves at all energy levels; **Peak Gained** Non-monotonic at higher energy levels only; **Peak Lost** Non-monotonic at low energy levels only; **All Non Monotonic** Non-monotonic SR curves at all energy levels. The Peak Gained and Peak Lost classes are so called as they are characterised by an appearing or disappearing local maxima (peak) as energy levels increase. Classes grouped by rows have similar high-energy behaviour (100 ATP); classes grouped by columns have similar low-energy behaviour (0.1 ATP).

Third, parameters which are nominally similar in function can have different, sometimes opposite effects on the system. For example, consider the forward phosphorylation reaction rates *κ*_f2_ (row 2, column 3) and *κ*_f3_ (row 2, column 4). These reaction rates both both determine how quickly the second phosphoryl group is ‘promoted’ from binding site 2 to 3 in the HK, but for different initial HK states (HK3 and HK5 respectively). According to Fig. 9a, these rates influence whether the SR curve is monotonic at low energy levels. Paradoxically, they exhibit opposite trends: If *κ*_f2_ is low, non-monotonic (blue) and peak-lost (red) solutions dominate. Both of these classes are non-monotonic at low energy levels; In contrast, if *κ*_f3_ is low, all-monotonic (green) and peak-gained (orange) solutions dominate, which demonstrate monotonicity at low energy.

We can perform a similar analysis for the peak-shift classes (Fig. 9b). This time, when the initial HK phosphorylation rate *κ*_ap1_ is low, the SR curve peak tends to move to the right as energy increases; that is, more stimulus is required to maximise the response when energy is abundant. Strangely, the converse is true for *κ*_a1_ (row 1, column 1), the rate of signal activation for the same pathway. *κ*_f2_ and *κ*_f3_ again exhibit similarly mirrored behaviour as before.

More investigation is required into how these parameters influence the SR curve, and which parameter combinations determine the unorthodox TCS signalling behaviour.

## Discussion

### Key insights from the TCS model

Here we have taken an energy-based approach to modelling TCSs and we have shown how this thermodynamic perspective leads to new and unexpected insights:

1. Energy limits the maximum signal expression by the response curve.
2. Energy affects the TCS temporal response curve sensitivity.
3. Noisy signals are sensitive to energy, but some systems are still able to filter out noise effectively
4. Unorthodox TCSs can detect for specific stimulus levels; this behaviour is energy dependent.

We find that energy availability affects the signal strength of a TCS (Fig 3). This has implications for how well a cell can function in challenging, low energy environments. A lower signal response from an equivalent stimulus level means that the cell will not respond as “confidently” or rapidly as it should. A signal that is too weak to be detected may result in a false negative stimulus detection. This may not be detrimental in some cases, as the activities of the TCSs in a cell could be triaged according to need (for example, TCSs related to cell structure or metabolism may be prioritised [29]).

We find that the temporal signal sensitivity of a TCS increases as available energy decreases, particularly for orthodox TCSs (Fig 4). This implies that for some TCSs the signalling mechanism will become more selective and behave more like a biological switch as ATP levels are reduced. A more sensitive system takes longer to initially accept a signal, but will produce a stronger, more pronounced response once the threshold has been reached. Such a system will be prone to less false positives, and will more likely respond to a true stimulus. This response behaviour could be useful for a system which triggers energy-intensive operations within the cell. Bacteria have been shown to adjust protein abundance in starvation or other hostile conditions [29]. In low energy environments, where food is scarce or energy must be rationed carefully, a cell may not want to waste energy following a false positive signal for food.

Our models demonstrate that unorthodox TCSs tend to filter noisy stimuli better than orthodox TCSs (Fig 5–7). The signal variance and the mean signal reduction are lower for the unorthodox variants than for the orthodox variants. We also see that, while higher energy levels amplify noise in the orthodox system, the unorthodox system is more robust to this noise amplification. Scenarios where accurate signal detection is important may justify the extra operational cost of unorthodox TCSs. In *E. coli,* for example, the unorthodox TCS responsible for anaerobic metabolism TorS/TorR activates in the absence of oxygen (Table 3, see also [12]). An accurate signal is important for the cell so as to not trigger aerobic energy pathways without oxygen, and vice-versa.

This filtering feature is due to the structural differences between the variants. For the orthodox system, there are only three reactions that separate the input signal from the output response (Table 4). The noise amplitude and variance do not change noticeably; so all the noise is passed on to the output signal (Fig 7, top rows). For the unorthodox system, there is a sequence of steps, so rapid changes in input do not cause a major increase in flux across the system (Table 7). Our hypothesis is that the multiple steps delay the signal propagation enough so that random fluctuations do not have time to pass through the system. As a result, flux in the intermediate reactions is relatively smaller, and the output signal is less noisy (Fig 7, bottom rows). This noise-reducing property of the unorthodox TCS aligns with previous modelling studies [14,15].

How well a system can filter noise appears to be energy dependent. More energy in the system means that there is more activity (i.e. higher reaction flux) in intermediate reactions. With additional ATP, noisy variations in the signal are processed and propagated through much more quickly, ultimately leading to noisier outputs (Fig 6). The unorthodox system, with its multiple signalling pathways, is able to mitigate this noise even in high-energy contexts (Fig 7, bottom rows). To our knowledge, no other study has investigated this effect.

An interesting observation in the unorthodox TCS models is the appearance of peaks in the signal response curves (Fig 8); this effectively means that such systems can act as band-pass filters [30]. The maximum response expression level is not found when the signal is maximised; rather it is found at an intermediate stimulus value. An optimal response by the cell will involve moving towards this intermediate stimulus level (or alternatively, a strong response to move away from this level). We therefore suggest that these signalling systems are designed to select for a particular signal expression level. This behaviour is useful for a cell if, for example, the TCS is detecting a quantity such as pH via the concentration of H^+^ ions [3]. In some cases, it may not make sense to search for “minimum” or “maximum” pH, rather a particular pH band would be desirable. This non-monotonic signal exists in similar biological contexts, such as TCS models extended with redox regulation [18], or drug dose-response curves in pharmacology [31]. It is of interest to investigate how different signalling structures give to rise similar non-monotonic behaviours in the future.

Furthermore, some unorthodox TCS exhibit the unusual property of a disappearing signalling peak as energy levels changed (Fig 10. This means that in high (or low) energy environments, the solution behaves monotonically, where a ‘switch-like’ decision is required; but in low (or high) energy environments, the solution is fine-tuned to a desired stimulus level, as above. In other words, at one energy level the cell selects for a particular stimulus, but at different energy level a clear decision is made once a stimulus threshold is crossed.

We found that in some cases, peaks in the SR curve can shift horizontally with a change in energy (Fig 11). Much like a radio tuner honing in on a particular bandwidth, so too could such a TCS adjust the signal it is selecting for based on energy availability. This leads to greater adaptability available to a cell than previously thought. In the pH example, it may be the case that the cell reacts to different pH under different energy levels. Cells can adapt to hostile environments [29]; perhaps in low nutrient environments the cell seeks out slightly more acidic or basic conditions compared to an environment abundant in nutrient.

**Fig 11.**
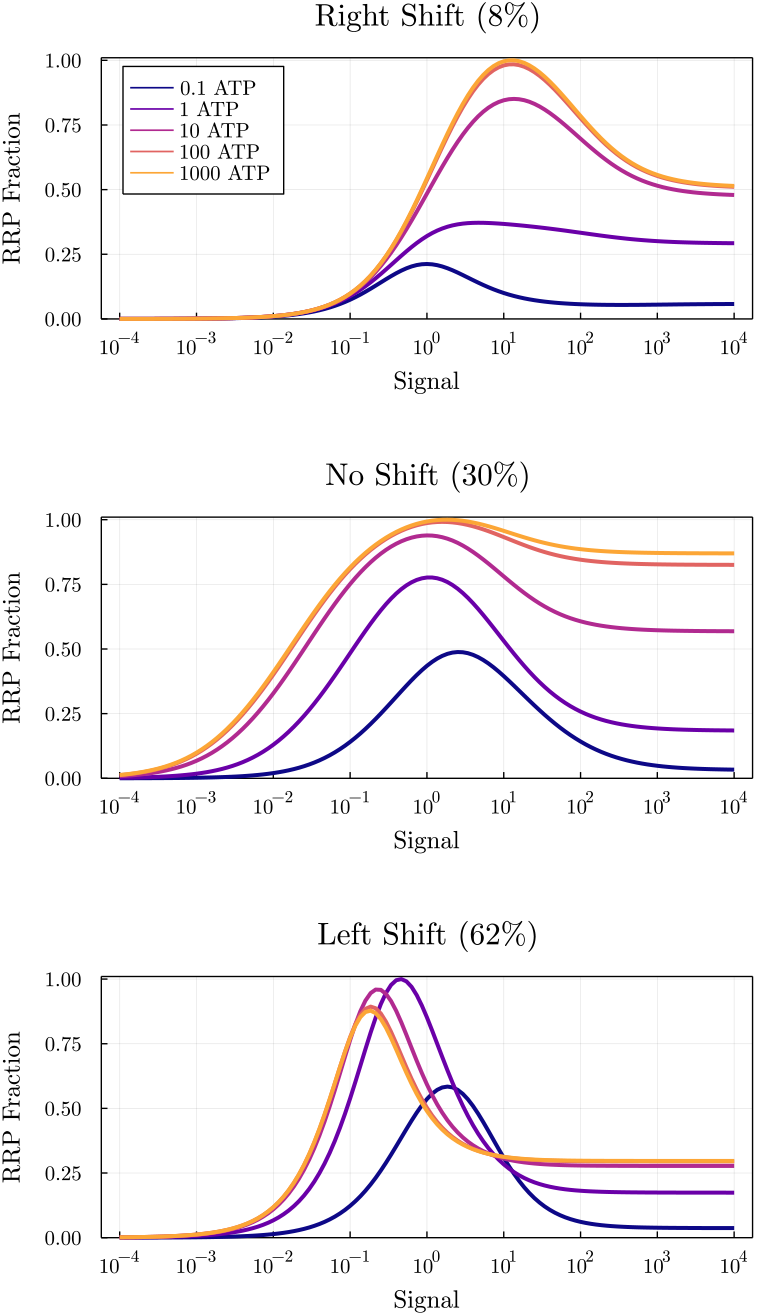
Representative examples of the SR peak-shift classes for the unorthodox TCS simulations. The proportions of each class are included in parentheses. Classes are only defined for the All Non Monotonic SR curve shapes, as these classes have a local maxima (peak) at all energy levels. These classes partition the SR curves based on how the horizontal position of the peak shifts as energy levels increase. This means that the signal that produces the maximum response expression level changes as a function of the energy level. **Right Shift** peak moves to the right; **Left Shift** peak moves to the left; **No Shift** no consistent or discernible movement in the peak (some movement is still permitted). Note that vertical shift also occurs alongside horizontal shift, and this may not be monotonic itself, such as in the Left Shift example.

These exotic behaviours are not rare occurrences in our analyses (Table 1). We can thus be reasonably confident that in some bacteria unorthodox TCSs exist whose signalling behaviour is non-monotonic. ATP-modulating SR curves are also not uncommon. This implies that for some cells the signalling behaviour could be adapted to new environments depending on the availability of ATP. These features highlight the wide array of adaptable signalling measures at the disposal of a cell, all within unorthodox TCS variants.

While a full study of a causal link between parameter selection and SR curve behaviour is beyond the scope of this investigation, our analysis provides some interesting insights (Fig 9a): all the thermodynamics constants of the species (the *K*’s) for each class follow the same distribution. The differences in parameter distributions only appear for the reaction rates (the *κ*’s). We see then that the SR behaviour of the TCSs is determined not by species-specific parameters of the molecules involved, but by the reaction rate parameters of the reaction system. We also detect a biological connection between the model and parameter and the behaviour. The autophosphorylation reaction rate parameter *κ*_ap1_, which determines binding rate of ATP to an empty HK, greatly influences whether an unorthodox TCS will exhibit signal selecting behaviour. If this reaction occurs too quickly relative to the other reactions, signal selection is unlikely to occur.

From this parameter distribution analysis we can conclude that it is not an individual molecular species that determines unusual behaviour; rather, it is the system-wide behaviour that matters. This supports the idea that a systems biology approach to modelling is required to understand the mechanisms; studying the structure of individual species alone will not be enough.

### ATP and Energy Modelling

Our study focuses on the energy requirements for TCS in bacteria. ATP is the ubiquitous source of chemical energy in biochemical reactions, so we have used the concentration of ATP as our proxy measure of energy availability for TCS. We run our simulations across an ATP gradient from ‘starvation’ to ‘abundance’. That is, when ATP levels are low, we interpret this as the bacteria being starved of a food source. Even in starvation or hostile conditions, cells can still continue to operate as normal [29].

The exact concentration of ATP can fluctuate greatly within a single population, even if the average concentration is stable. Yaginuma et al. [32] show that the distribution of ATP in *E. coli* follows a heavy-tailed right-skewed distribution. While the average concentration was 1.54mM, the actual values ranged from 14mM to 0mM (or negligible amounts). They theorise that “the diversity in the level of ATP may benefit the population by enabling individual cells to adopt different strategies under severe conditions” [32].

An alternative interpretation of “starvation conditions” is to imagine the amount of ATP allocated to the particular TCS in triage; certain signalling systems may not be as important in these scenarios, such as TCSs used to determine whether the cell should divide. Under this interpretation we are really modelling the concentration of local ATP allocated or available for a particular TCS system in a particular scenario. As some of our results show (e.g. Figs. 10–11), there may be additionally benefits to the cell when less ATP is allocated to some TCS.

### Synthetic biology

We would ultimately like to link TCS structural properties (e.g. size and complexity) to their function or purpose (e.g. temperature sensing or cell motility). The range of signalling behaviour and the adaptability of the TCS provides us with much scope for synthetic biology design [17, 20]. There are many parameters available in the design of a TCS to control the behaviour of the system (9 for orthodox, 35 for unorthodox). It may be possible that by adjusting the parameter value for a single component we can select which kind of behaviour we want in our cell: adaptable sensitivity, energy modulation, noise filtering, fine-tuned signal selection, signal peak formation, and signal peak tuning. Further investigation is needed to determine precisely how parameter choices cause these TCS properties. It is likely that underlying meta-parameters (e.g. the ratio of reaction rates) determine TCS behaviour and that we can use qualitative approaches to help in delineating different regimes of TCS behaviour [33].

Reaction rate parameters are notoriously difficult to determine from experimental observations [34] and depend on environmental conditions such as temperature and pressure. Thermodynamic constants, by contrast, are connected with the molecular species and can be transferred across to other mathematical models. It is possible to derive reaction rates from kinetic rates found in literature, provided some parameter values are known already (e.g. [35, 36]). Future work will focus on measuring reaction rates to determine whether unorthodox TCSs have observable signalling peaks or shifts in peaks.

Our study of energy-dependent effects of TCS in bacteria reveal novel insights into the capabilities of cellular signalling systems. These findings generate new functional hypotheses that are experimentally testable [37]. We suggest that future experiments in TCS should investigate how signalling systems behave in low-energy environments; how signalling systems filter out noise; and how unorthodox TCS adapt their signalling behaviour.

## Methods

We have modelled TCS dynamics with a system of ODEs [1,4,7,10,11,14–16]. Our chemical reaction system notation is based on the model described by Kim and Cho [15]. Unlike their study, we explicitly retain ATP in the autophosphorylation chemical reactions. Note that the stimulus *S* is a generic input that may not be chemical (e.g. light, heat, envelope stress). We have treated S as such a stimulant which is ‘consumed’ by the signalling pathway. If the stimulant were a chemical species, the model would need to be modified so that the stimulant is regenerated during the cycle to ensure mass balance; we do not consider this case here. The full models for the orthodox and unorthodox TCS, the chemical reaction networks, ODE models, and parameter definitions, can be found in Appendix B in S1 Text.

ATP enters the system through the autophosphorylation of HK (Table 2). Though included in the reaction system, this is usually dropped from the model, since it is not considered rate limiting [14,15]. Instead of assuming an unlimited concentration of ATP, we assume a finite (yet constant) supply of ATP. In order to maintain mass and energy conservation laws, the reaction system includes ADP and Pi (inorganic phosphate group) as well. For thermodynamic feasibility, we also require all chemical reactions to be reversible.

**Table 2.** Orthodox and Unorthodox TCS Chemical Reaction Systems. Reaction systems for the TCS models used in this study, based on those used in Kim and Cho [15]. Parallels exist between the reaction types of the orthodox and unorthodox variants, except for the forward phosphorelay, which only applies to the latter.

| Reaction Name | Orthodox TCS Reactions | Unorthodox TCS Reactions |
| --- | --- | --- |
| Signal Activation | $\text{HK} + \text{S} \xrightleftharpoons{\kappa_a} \text{HKS}$ | $\text{HK1} + \text{S} \xrightleftharpoons{\kappa_{a1}} \text{HK1S}$<br>$\text{HK3} + \text{S} \xrightleftharpoons{\kappa_{a2}} \text{HK3S}$<br>$\text{HK4} + \text{S} \xrightleftharpoons{\kappa_{a3}} \text{HK4S}$<br>$\text{HK7} + \text{S} \xrightleftharpoons{\kappa_{a4}} \text{HK7S}$ |
| Autophosphorylation | $\text{HKS} + \text{ATP} \xrightleftharpoons{\kappa_{ap}} \text{HKP} + \text{ADP}$ | $\text{HK1S} + \text{ATP} \xrightleftharpoons{\kappa_{ap1}} \text{HK2} + \text{ADP}$<br>$\text{HK3S} + \text{ATP} \xrightleftharpoons{\kappa_{ap2}} \text{HK5} + \text{ADP}$<br>$\text{HK4S} + \text{ATP} \xrightleftharpoons{\kappa_{ap3}} \text{HK6} + \text{ADP}$<br>$\text{HK7S} + \text{ATP} \xrightleftharpoons{\kappa_{ap4}} \text{HK8} + \text{ADP}$ |
| Forward Phosphorelay | *n/a* | $\text{HK2} \xrightleftharpoons{\kappa_{f1}} \text{HK3}$<br>$\text{HK3} \xrightleftharpoons{\kappa_{f2}} \text{HK4}$<br>$\text{HK5} \xrightleftharpoons{\kappa_{f3}} \text{HK6}$<br>$\text{HK6} \xrightleftharpoons{\kappa_{f4}} \text{HK7}$ |
| Phosphotransfer | $\text{HKP} + \text{RR} \xrightleftharpoons{\kappa_{pt}} \text{HK} + \text{RRP}$ | $\text{HK4} + \text{RR} \xrightleftharpoons{\kappa_{pt1}} \text{HK1} + \text{RRP}$<br>$\text{HK6} + \text{RR} \xrightleftharpoons{\kappa_{pt2}} \text{HK2} + \text{RRP}$<br>$\text{HK7} + \text{RR} \xrightleftharpoons{\kappa_{pt3}} \text{HK3} + \text{RRP}$<br>$\text{HK8} + \text{RR} \xrightleftharpoons{\kappa_{pt4}} \text{HK5} + \text{RRP}$ |
| Degradation | $\text{RRP} \xrightleftharpoons{\kappa_d} \text{RR} + \text{Pi}$ | $\text{HK8} \xrightleftharpoons{\kappa_{d1}} \text{HK6} + \text{Pi}$<br>$\text{HK5} \xrightleftharpoons{\kappa_{d2}} \text{HK2} + \text{Pi}$<br>$\text{HK3} \xrightleftharpoons{\kappa_{d3}} \text{HK1} + \text{Pi}$<br>$\text{HK7} \xrightleftharpoons{\kappa_{d4}} \text{HK4} + \text{Pi}$<br>$\text{RRP} \xrightleftharpoons{\kappa_{d5}} \text{RR} + \text{Pi}$ |

### Bond Graphs

We represent our TCS chemical reaction networks using biochemical bond graphs. For an introduction on bond graphs, see [38, 39]; for applications in biochemistry, see [23,24,35,36,40].

Bond graphs represent physical systems as mathematical graphs, with nodes representing either system elements (*components*) or conservation laws, (*junctions*), and edges representing flows of energy, (*bonds*).

We can represent any chemical reaction network as a bond graph network. Chemical species are represented by one-port **C** components that track concentrations; reactions are represented by two-port **Re** components, where the two ports correspond to the reactants and products of the reaction (Note: since all reactions are reversible, the distinction between reactants and products may not always apply). Species which appear in multiple reactions connect to a **0**-junction; these junctions conserve the mass of a species. Species on the same side of a reaction (the reactants or products) are combined with a **1**-junction; these junctions conserve chemical potential *μ* and represent the combined potential energy for the reaction.

As an illustrative example, take the combined signal activation and autophosphorylation reactions for the orthodox TCS (Table 2).

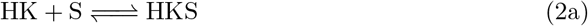

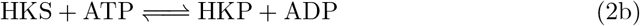

This reaction subsystem is represented as a bond graph in Fig. 12A. HK, HKS, HKP, ATP, ADP, and S each have a **C** component that stores the chemical potential energy of each species. **Re:Activate** and **Re:Autophos** are the reaction events in the system. These spend chemical potential energy and describe flow by a constitutive relation (see Model Derivation). HK and S combine to form HKS, so they are connected with a **1**-junction. HKS appears in both reactions and so is split with a **0**-junction. Each chemical species appears only once in order to ensure they are conserved across all reactions in the system.

**Fig 12.**
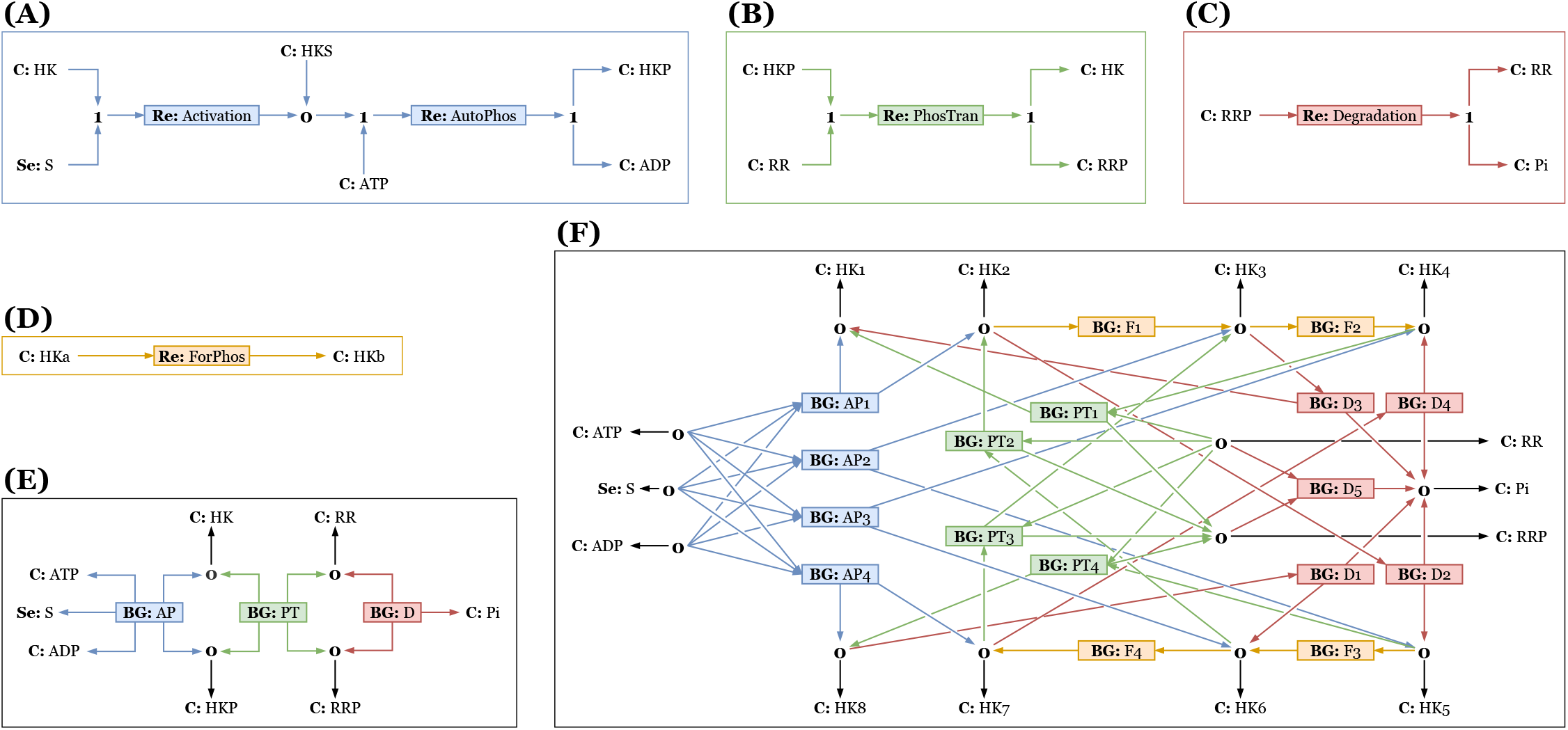
Bond graph representations of all TCS reactions and models: **(A)** Signal Activation and Autophosphorylation **(B)** Phosphotransfer **(C)** Degradation **(D)** Forward Phosphorelay **(E)** Orthodox TCS **(F)** Unorthodox TCS. Bond graphs **(A)-(D)** form the components used in the hierarchical TCS models **(E)-(F)**.

This structure can be iteratively generated for our TCS models from Table 2. Figs. 12A-D show the bond graphs for each reaction subsystem. These graphs become **BG** components used to build our hierarchical TCS models Figs. 12E-F. Species in the lower level bond graphs which appear in other bond graphs are exposed to the higher level. The chemostat species ATP, ADP, Pi, and S are all held at constant concentrations. Note that the unorthodox TCS model uses multiple instances of the same reaction submodel. For this study we generated our bond graph models using the Python package BondGraphTools [41].

### Model Derivation

Energy flow is described using the generalised units of *effort* [Joules/x] and *flow* [x/second], where x is a tracked quantity for a particular physical system. For example, if x is the charge *q* in Coulombs, then the effort is [Joules/*x*] ≡ [Volts] and the flow is [q/second] ≡ [Amperes]. The relationship between the effort and flow in a component (e.g. an electrical resistor) is described by a constitutive relation (e.g. *V* = *IR*). In addition to constitutive relations we have conservation laws that conserve mass, momentum, charge, or a particular species (e.g. Kirchhoff’s Current Law).

For biochemical bond graphs, the effort variable is the chemical potential μ [J/mol].

Each reaction species *i* has its own chemical potential, defined by

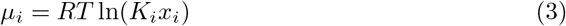

where *R* [J/mol/K] is the universal gas constant, *T* [K] is the absolute temperature, *K_i_* [mol^-1^] is the species specific thermodynamic constant, and *x_i_* [mol] is the concentration of species *i*.

The flow variable is the reaction flux *v* [mol/s], which is the reaction rate for each reaction in the model. Assuming mass action kinetics, the flux through each reaction *j* is defined as

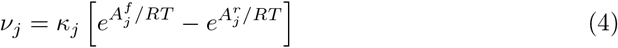

where *κ_j_* [mol/s] is the molar reaction rate for reaction *j*, and 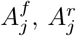 [J/mol] are the sums of the chemical potentials of the reactants (forward affinity) or products (reverse affinity), respectively.

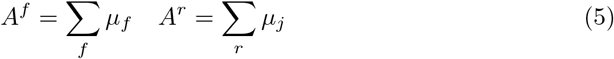

where *f* and *r* are the forward and reverse reactants, respectively.

To derive mass action kinetics, we can substitute Eq 3 and Eq 5 into Eq 4:

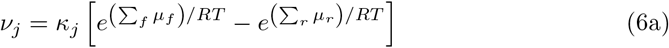

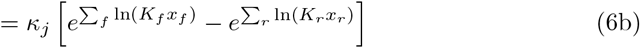

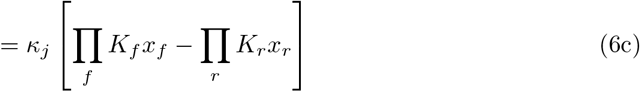

When Eq 6c is expanded, the reaction rates Kj and species’ thermodynamic constants *K_i_* can be combined to form the familiar kinetic rate constants of mass action kinetics.

Note that *κ* determines only the rate of a reaction, not the direction. The reaction flux direction is determined by whichever is greater of the product of *K_f_ x_f_* or *K_r_x_r_*. The species constants and concentrations themselves determine the reaction direction. This is in contract to standard mass action kinetic models where the forward and backward reaction rates can be selected independently from one another.

The concentration of each species over time is

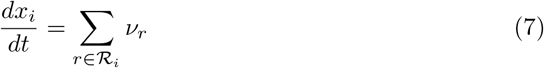

where 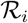 is the set of reactions that involve species *x_i_*. With these equations and the bond graph structure described in Fig 12 we can algorithmically generate a system of nonlinear ODEs. This method is used to generate our TCS ODE models (Appendix B, S1 Text).

### Parameter Selection

In order to explore the family of behaviours that observed in TCS we have performed a parameter sensitivity study on the model parameters. We pre-generated 10,000 parameter sets for each model in order to capture likely (normalised) biological parameter ranges. These parameters sets are reused for each model when comparing similar simulations across ATP concentrations or other control variable.

For convenience, all initial conditions, parameters, and solution values used in our simulations are normalised and/or chosen as abstract quantities rather than experimentally recorded values. The universal gas constant *R* and temperature *T* are both normalised so that *RT* = 1 in Eq 3 and Eq 4. For our purposes, the use of abstract parameters is sufficient to compare the overall structure and behaviour of the two models under different conditions.

The sampling ranges for the parameters in the models can be found in Table 5 and Table 7. The thermodynamic constants K for the core component species (HK, RR, and their variants) are sampled on a log_10_ scale from 0.1 to 10. The reaction rates *κ* are sampled in the same way:

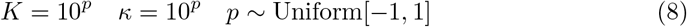

Although exact parameter ranges for most TCS parameters are unknown, we can still use some physical and biological intuition to constrain our models. This limits the parameter space to exclude physiologically unrealistic parameter combinations. This helps refine our parameter search by filtering out the unrealistic or trivial TCS parameter sets, such as those which would flow backwards or not function. We do this with two changes.

First, due to the function of ATP as a reaction activator, we know that it will have a larger chemical potential than any of the other species involved. Likewise, ADP and Pi do not themselves drive reactions forward, and so will have relatively lower chemical potentials. If this were not the case, then we would find our reactions primarily travelling in the opposite direction. Since we are dealing in primarily abstract quantities, we only focus on the relative magnitudes of parameters. Therefore, in our models we have used *K*_ATP_ = 100, *K*_ADP_ = 0.1, and *K*_Pi_ = 0.001.

Second, we know *a priori* the dominant direction of all reactions. These are the reactions presented as irreversible in earlier TCS studies. We do not want to set all the reactions to be irreversible, as that would break the thermodynamic requirements and limit reaction possibilities. We can instead sample chemical potentials so that forward reactions are *favoured* rather than eliminating reverse reactions.

We have achieved this by ordering the sampled thermodynamic constants *K* before assigning them to specific species. In other words, for a model with *N* species, we sample *N* parameters, sort in descending order, and assign to the species in order of appearance in the reaction system (Eq 9).

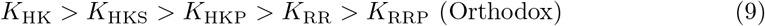

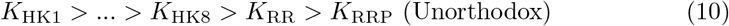

This method does not mean all the reactions are now irreversible; rather it biases the reactions in a meaningful direction, while still allowing reverse reactions to occur. Note that there are multiple ways to order the unorthodox HK states (HK1...HK8), since their order of appearance in the system is not strictly linear. Nevertheless, this is a fairly natural ordering that achieves the desired effect.

### Initial Conditions

The initial conditions for the core TCS species in all simulations are [RR]_0_ = 10, [HK]_0_ = 1 (or [HK1]_0_ = 1 for the unorthodox model), and [HK*x*] = 0 for all other HK states *x*. Likewise, we set [ADP]_0_ = [Pi]_0_ = 0.

We used five levels of available ATP concentration in our model:

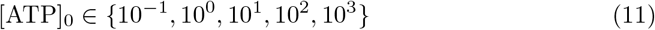

This gave us an even spread of energy levels across a log10 scale.

For the first set of simulations (Figs. 3, 4) the stimulus input function was set to a constant value *S*(*t*) = *S*_0_ = 1. For the SR curve simulations (Figs. 8, 10, 11) we used 121,000 log_10_-spaced values of *S*_0_ ∈ [10^-4^,10^4^].

When modelling a noisy stimulus (Figs. 5, 6, 7), we generate the noise with a piece-wise function *s_k_*(*t*) defined by:

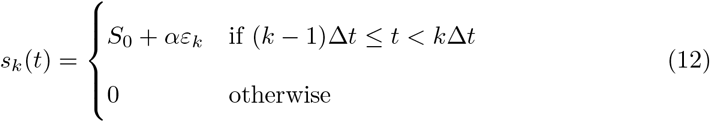

where the noise 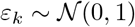 is distributed normally, *α* = 0.5 is the magnitude of the noise, *S*_0_ = 1 is the constant baseline, and Δ*t* = 1 is the length of interval *k.* We then define the stimulus input function as:

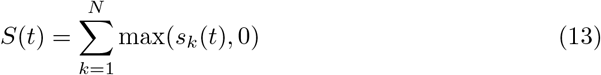

where *N* = 200 is the total number of intervals. We include the max function in the summation so that the input is always positive. The noise terms *ε_k_* are all pregenerated so that they can be reused and direct comparisons made across all simulations.

### Simulation

For each TCS model we ran 10,000 simulations with pregenerated randomly sampled parameters at the five determined ATP levels and a constant stimulus input (100,000 in total). For the noisy stimulus scenario, we ran 500 simulations per model per ATP level (5,000 in total). We solved our ODE systems numerically using the Differential Equations library in Julia [42]. The simulations were all run for *t* ∈ [0 200], which was found to be sufficient to capture all steady state behaviour for the parameter ranges studied.

Each simulation output included the RRP concentration over time (response curve). To aggregate the response curves we first normalise by the maximum response value (to get an RRP fraction between 0 and 1), and then take the average across all parameter sets for each curve group (grouped by model, ATP, and stimulant). This is done for example in the results in Fig. 3.

The noise filtering performance measures (Fig. 6) were calculated for *t* ∈ [100 200]. We used a generous warm-up period (around half the simulation time) to allow for the wide range of possible simulations to reach steady state.

## Supporting information

**S1 Text Supplementary material. Appendix A**: TCS species and functions found in *E. coli.* **Appendix B**: Chemical reactions, parameters, and ODE models of orthodox and unorthodox TCS. **Appendix C**: Additional figures.

## S1 Text

### A TCS in *E. coli*

**Table 3.** Two Component Systems (TCS) found in *E. coli,* with their recorded cellular functions. First table lists orthodox TCS; second table lists unorthodox TCS. The average amino acid length for the orthodox proteins are less than the corresponding unorthodox proteins. Source: EcoCyc Database [12]

| HK/RR | Amino Acid Number | Function |
| --- | --- | --- |
| AtoS/AtoC | 608/461 | Metabolism |
| BaeS/BaeR | 467/240 | Envelope homeostasis |
| BasS/BasR | 363/222 | Polymyxin resistance |
| BtsS/BtsR | 561/239 | Pyruvate sensing |
| CheA/CheB | 654/349 | Chemotaxis |
| CheA/CheY | 654/129 | Chemotaxis |
| CpxA/CpxR | 457/232 | Envelope homeostasis |
| CreC/CreB | 474/229 | Regulates aerobic growth |
| CusS/CusR | 480/227 | Copper response |
| DcuS/DcuR | 543/239 | Dicarboxylates response |
| DpiB/DpiA | 552/226 | Citrate fermentation |
| EnvZ/OmpR | 450/239 | Osmolarity sensing |
| GlrK/GlrR | 475/444 | Envelope homeostasis |
| HprS/HprR | 452/223 | Hydrogen peroxide sensitivity |
| KdpD/KdpE | 894/225 | Potassium transport |
| NarQ/NarP | 566/215 | Response to nitrite |
| NarX/NarL | 598/216 | Response to nitrite |
| NtrB/NtrC | 349/470 | Nitrogen response |
| PhoQ/PhoP | 486/223 | Response to magnesium |
| PhoR/PhoB | 431/229 | Phosphohate regulation |
| PyrS/PyrR | 565/244 | Pyruvate sensing |
| QseC/QseB | 449/219 | Cell motility |
| RcsD/RcsB | 890/216 | Envelope homeostasis |
| RstB/RstA | 433/239 | Acid tolerance |
| UhpB/UhpA | 500/196 | Metabolism |
| ZraS/ZraR | 465/441 | Envelope homeostasis |
| **Mean** | **531/263** |  |
| ArcB/ArcA | 778/238 | Respiratory control |
| BarA/UvrY | 918/218 | Hydrogen peroxide sensitivity |
| EvgS/EvgA | 1197/204 | Drug resistance |
| TorS/TorR | 915/230 | Anaerobic energy production |
| **Mean** | **952/223** |  |

### B TCS Models

#### B.1 Orthodox TCS

##### Chemical Reactions

**Table 4.**
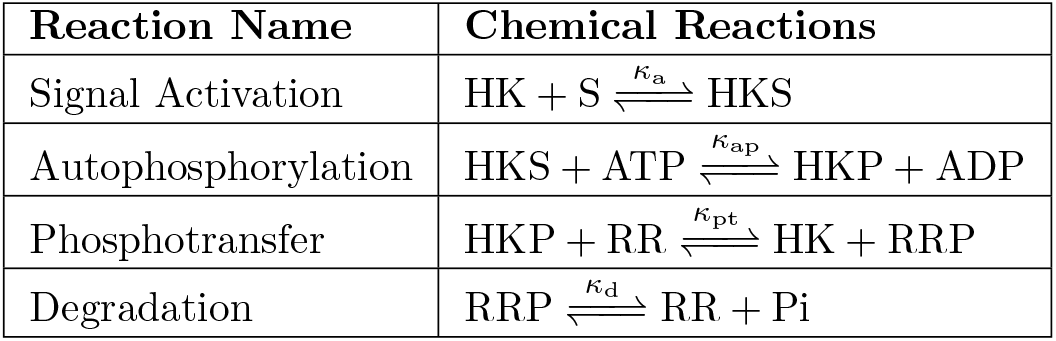
Chemical reaction system for our orthodox TCS model. This system is an extension on the model used by Kim and Cho [15]. Here we explicitly include ATP in the reaction system with a two-step phosphorylation of HK, and allow reactions to be reversible. Note that the input stimulant S may be an intangible quantity such as heat or mechanical stress, and so is not conserved in the model.

##### Parameters

**Table 5.**
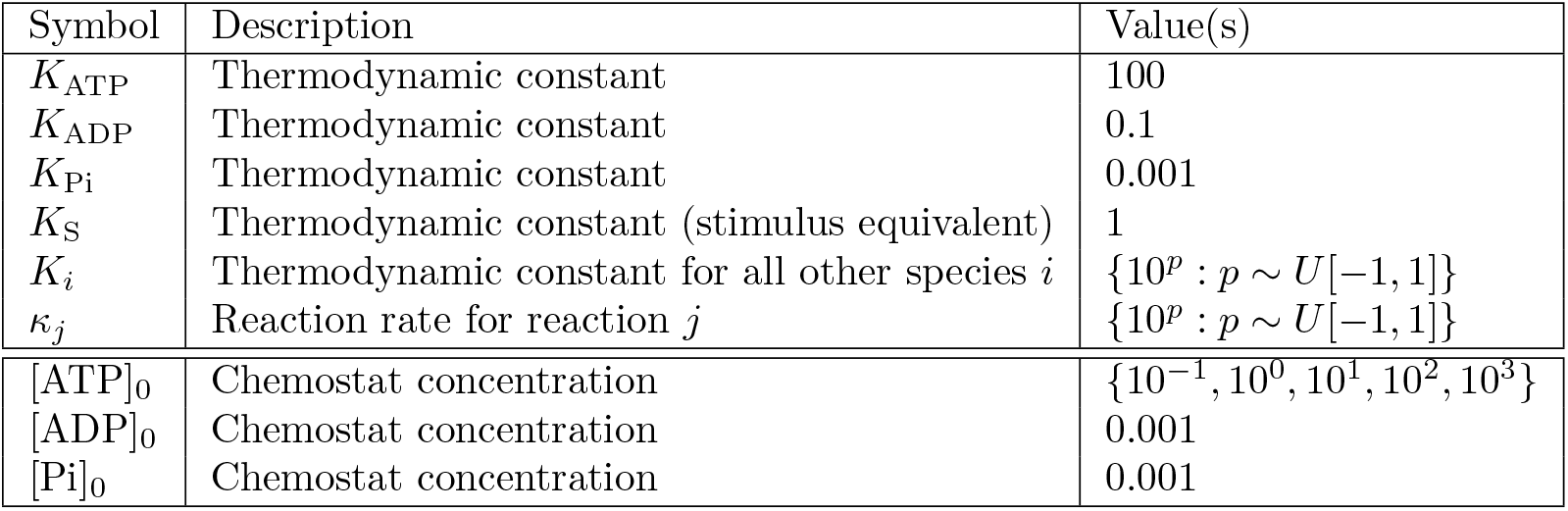
ODE model parameters for the orthodox TCS.

##### ODE Model

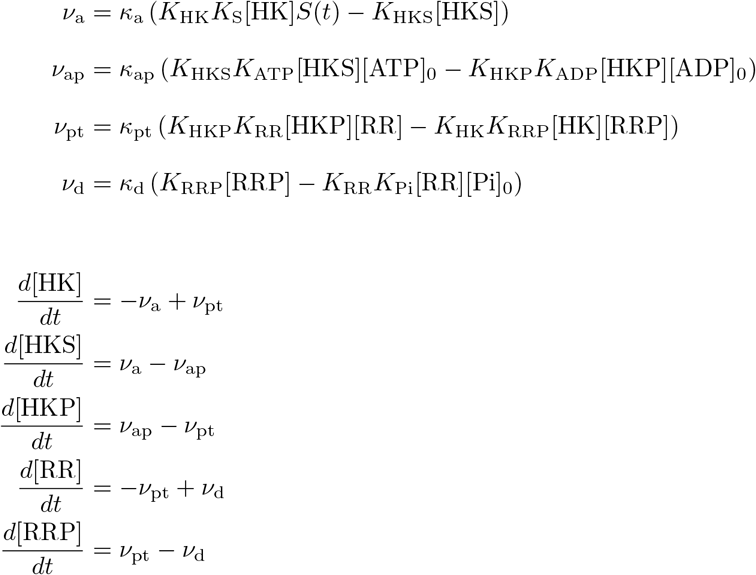

#### B.2 Unorthodox TCS

##### Chemical Reactions

**Table 6.** Chemical reaction system for our unorthodox TCS model. This system is an extension on the model used by Kim and Cho [15]. Here we explicitly include ATP in the reaction system with a two-step phosphorylation of HK, and allow reactions to be reversible. Note that the input stimulant S may be an intangible quantity such as heat or mechanical stress, and so is not conserved in the model.

| Reaction Name | Chemical Reactions |
| --- | --- |
| Signal Activation | $\text{HK1} + \text{S} \xrightleftharpoons{\kappa_{a1}} \text{HK1S}$<br>$\text{HK3} + \text{S} \xrightleftharpoons{\kappa_{a2}} \text{HK3S}$<br>$\text{HK4} + \text{S} \xrightleftharpoons{\kappa_{a3}} \text{HK4S}$<br>$\text{HK7} + \text{S} \xrightleftharpoons{\kappa_{a4}} \text{HK7S}$ |
| Autophosphorylation | $\text{HK1S} + \text{ATP} \xrightleftharpoons{\kappa_{ap1}} \text{HK2} + \text{ADP}$<br>$\text{HK3S} + \text{ATP} \xrightleftharpoons{\kappa_{ap2}} \text{HK5} + \text{ADP}$<br>$\text{HK4S} + \text{ATP} \xrightleftharpoons{\kappa_{ap3}} \text{HK6} + \text{ADP}$<br>$\text{HK7S} + \text{ATP} \xrightleftharpoons{\kappa_{ap4}} \text{HK8} + \text{ADP}$ |
| Forward Phosphorelay | $\text{HK2} \xrightleftharpoons{\kappa_{f1}} \text{HK3}$<br>$\text{HK3} \xrightleftharpoons{\kappa_{f2}} \text{HK4}$<br>$\text{HK5} \xrightleftharpoons{\kappa_{f3}} \text{HK6}$<br>$\text{HK6} \xrightleftharpoons{\kappa_{f4}} \text{HK7}$ |
| Phosphotransfer | $\text{HK4} + \text{RR} \xrightleftharpoons{\kappa_{pt1}} \text{HK1} + \text{RRP}$<br>$\text{HK6} + \text{RR} \xrightleftharpoons{\kappa_{pt2}} \text{HK2} + \text{RRP}$<br>$\text{HK7} + \text{RR} \xrightleftharpoons{\kappa_{pt3}} \text{HK3} + \text{RRP}$<br>$\text{HK8} + \text{RR} \xrightleftharpoons{\kappa_{pt4}} \text{HK5} + \text{RRP}$ |
| Degradation | $\text{HK8} \xrightleftharpoons{\kappa_{d1}} \text{HK6} + \text{Pi}$<br>$\text{HK5} \xrightleftharpoons{\kappa_{d2}} \text{HK2} + \text{Pi}$<br>$\text{HK3} \xrightleftharpoons{\kappa_{d3}} \text{HK1} + \text{Pi}$<br>$\text{HK7} \xrightleftharpoons{\kappa_{d4}} \text{HK4} + \text{Pi}$<br>$\text{RRP} \xrightleftharpoons{\kappa_{d5}} \text{RR} + \text{Pi}$ |

##### Parameters

**Table 7.**
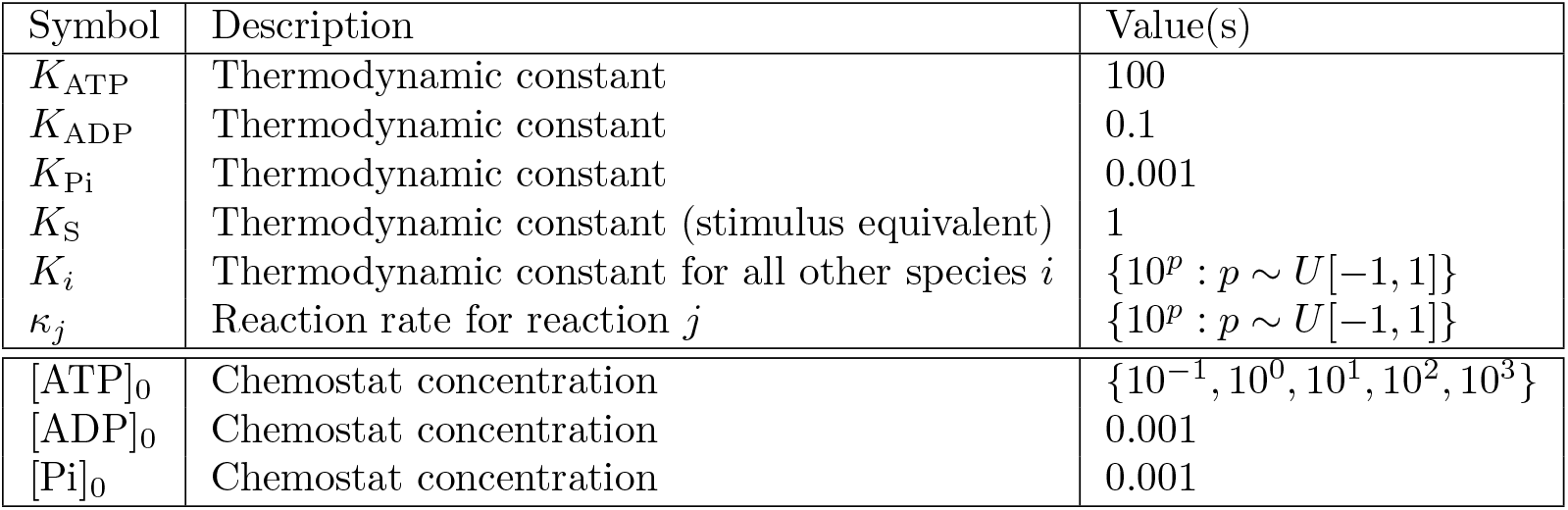
ODE model parameters for the unorthodox TCS.

##### ODE Model

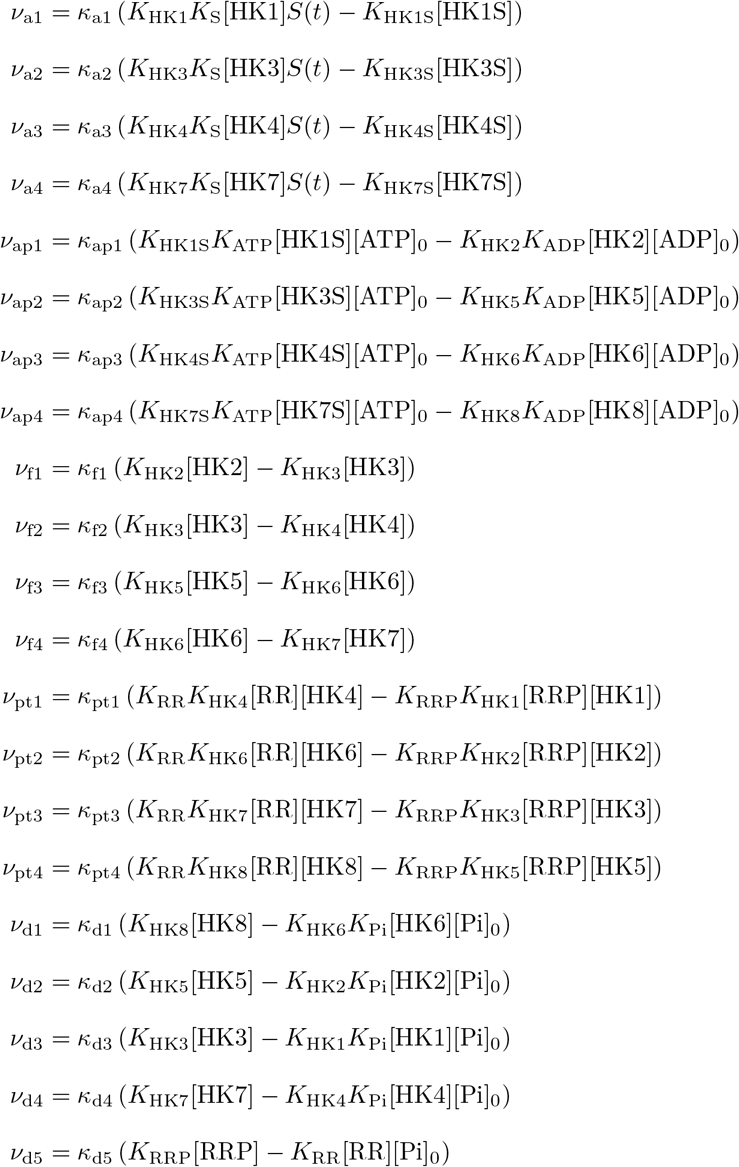

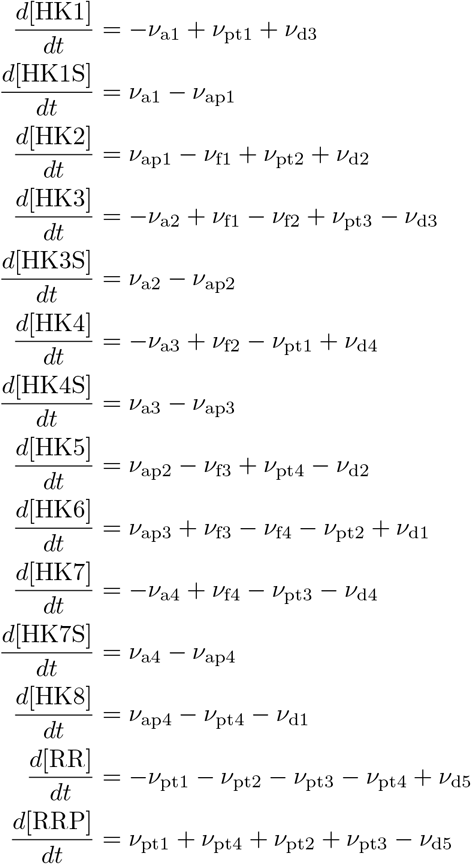

### C Additional Figures

**Fig 13.**
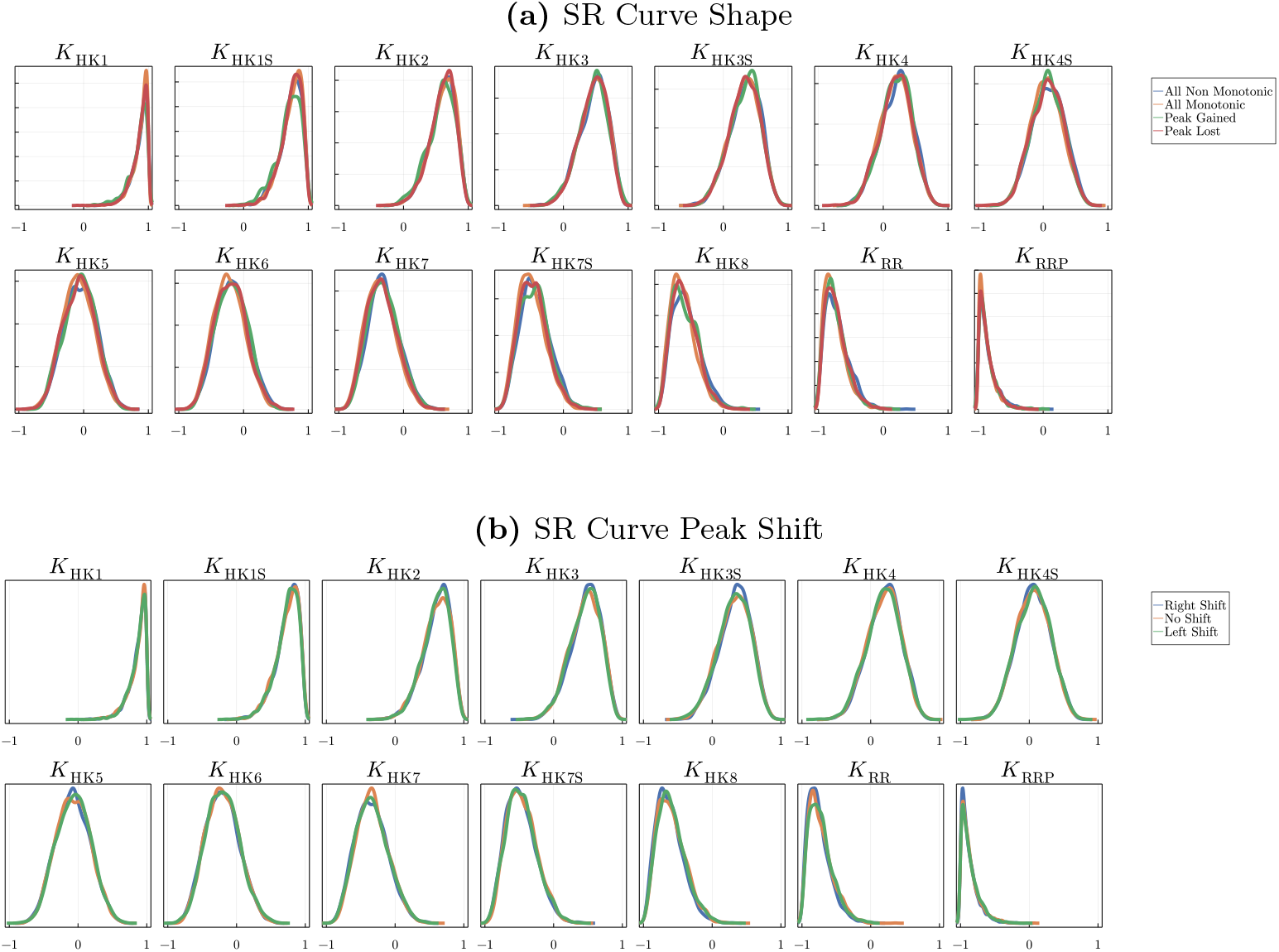
Log distribution of the sampled thermodynamic constants *K* used for the unorthodox TCS numerical simulations, split by **(a)** SR curve shape and **(b)** SR peak shift. The thermodynamic constants defined in Table 7 in Appendix B. Parameters are sampled from a log-uniform distribution (Eq. 8). The distributions for these parameters all follow roughly the same distribution, so we observe that these parameters do not affect SR shape or peak-shift classification (contrast with Fig. 9 in the main text). Note that these parameters follow a skewed normal distribution trend due to how the parameters are sampled; see Methods section in main text for more details.

